# Phage-encoded sigma factors alter bacterial dormancy

**DOI:** 10.1101/2021.11.12.468384

**Authors:** DA Schwartz, BK Lehmkuhl, JT Lennon

## Abstract

By entering a reversible state of reduced metabolic activity, dormant microorganisms are able to tolerate suboptimal conditions that would otherwise reduce their fitness. Dormancy may also benefit bacteria by serving as a refuge from parasitic infections. Here we focus on dormancy in the *Firmicutes*, where endospore development is transcriptionally regulated by the expression of sigma factors. A disruption of this process could influence the survivorship and reproduction of phages that infect spore-forming hosts with implications for coevolutionary dynamics. Here, we characterized the distribution and diversity of sigma factors in nearly 3,500 phage genomes. Homologs of sporulation-specific sigma factors were identified in phages that infect spore-forming hosts. Unlike sigma factors required for phage reproduction, the sporulation-like sigma factors were non-essential for lytic infection. However, when expressed in the spore-forming *Bacillus subtilis*, sigma factors from phages activated the bacterial sporulation gene network and reduced spore yield. Our findings suggest that the acquisition of host-like transcriptional regulators may allow phages to manipulate a complex and ancient trait in one of the most abundant cell types on Earth.

## INTRODUCTION

Dormancy is a life history strategy that allows individuals to enter a reversible state of reduced metabolic activity. An example of convergent evolution, it has independently arisen throughout the tree of life as a means of coping with fluctuating and unpredictable environments^1^. Dormancy is particularly prevalent among microbial life forms where it contributes to the persistence and fitness of populations in environments where variables like pH, oxygen, and resource availability are suboptimal for growth and reproduction^2^. In addition to buffering populations against abiotic features of the environment, dormancy may be reinforced through dynamics that arise from species interactions. For example, dormancy diminishes the strength of competition, which in turn can promote species coexistence^3^. In addition, dormancy may benefit populations by serving as a refuge against predator consumption or parasite infection^4,5^.

Among microorganisms, dormancy can protect hosts from viral parasites in a number of ways. As cells transition into an inactive state, they often undergo morphological changes that affect how viruses physically interact with their host. For example, the formation of dormant cells, such as cysts and spores, often involves the development of a thick exterior coating^6–8^ that masks the surface molecules used by viruses for attachment^9–12^. Even if a virus is able to gain entry into a dormant cell, parasite productivity will be low owing to constraints imposed by the host’s reduced metabolism^13–16^. Furthermore, viral defense genes are often located in proximity to genes that regulate dormancy and cell suicide, suggesting that dormancy may contribute to multilayered protection against viral infection^17,18^. For example, virus-induced dormancy has been linked to CRISPR-Cas systems in bacteria^20^ and archaea^19,20^. As a physiological refuge^21^, dormancy can confer herd immunity and diminish the spread of epidemics^22^, which may ultimately shape host-virus coevolutionary dynamics.

A take-home lesson from studies on antagonistic coevolution is that host defenses are prone to being overcome by viruses^23,24^. One general mode of virus adaptation involves the acquisition of host genes. Viral genomes commonly encode homologs of genes that are involved in host metabolism. These so-called “auxiliary” genes can alter cellular processes in ways that affect virus fitness^25^. Originally motivated by the discovery of photosynthesis genes in marine cyanophages^26,27^, auxiliary genes have been implicated in host nutrition (e.g., N and P metabolism) and energy acquisition (sulfur oxidation, fermentation, etc.), along with basic cellular functions such as protein translation and bacterial communication via quorum sensing^28–30^. In addition, some virus genomes contain host defense genes, which has led to speculation that auxiliary genes may modify parasite infectivity and reproduction^31,32^. Similarly, some phage genomes have been reported to have genes similar to those required for the development of endospore-forming bacteria ^33–42^. Phages might use sporulation homologs to inhibit their host from entering a dormancy refuge, thereby enhancing the reproductive component of parasite fitness. Alternatively, phages might exploit sporulation in a way that extends longevity and thereby enhance the survivorship component of fitness. This could happen through a process known as entrapment whereby a phage genome is translocated into the developing spore resulting in the production of a “virospore”^12,43–46^. Analogous to pseudolysogeny, the phage genome is protected by the endospore from conditions that would otherwise contribute to phage decay without it being integrated into the host chromosome^45^. When environmental conditions improve, the dormant cell undergoes germination, and the phage resumes its lytic reproductive cycle.

As a complex form of dormancy, sporulation presents phages with many opportunities for intervention. For proper development, sporulation requires the coordinated regulation of a large gene network^47–49^. The central regulatory module of sporulation relies on the activity of sigma factors, the exchangeable subunit of the transcriptional machinery which dictates promoter specificity of RNA polymerase^50^. Among bacteria, a primary sigma factor (*sigA* in *Bacillus subtilis)* is essential for growth, reproduction, and other housekeeping processes in a wide range of bacteria^51^. During *B. subtilis* spore development *sigA* is swapped out by a cascade of sporulation-specific sigma factors, each driving the expression of a subset of sporulation genes in distinct cellular compartments at specific times^52^. Following an asymmetrical cell division, gene expression in the maturing spore (i.e., forespore) is driven first by *sigF* and then *sigG*, while expression in the mother cell is driven by *sigE* and then *sigK*. Sigma factors are also encoded by some phage genomes, where they regulate phage gene expression during different stages of lytic development^52,53^. More recently, homologs of sporulation-specific sigma factors have been identified in phage genomes^34,41,42,54^, yet their function has not been explored. As central nodes of the sporulation gene network, sigma factors could be coopted by phages to modify the outcome of sporulation.

Here, we use a combination of bioinformatics and laboratory experiments to test whether homologs of sporulation-specific sigma factors can be used by phages to manipulate host dormancy. Using sequence homology and phylogenetic analyses, we identify and classify hundreds of phage-encoded sigma factors. We find that phages capable of infecting spore-forming hosts preferentially encode sigma factors that are homologous to the forespore-specific sigma factors, *sigF* and *sigG*. When homologs of sporulation-specific sigma factors are expressed in *B. subtilis*, we observed that conserved phage-encoded homologs alter host gene expression and reduce spore yield. Together, our findings have implications for understanding dormancy dynamics in environmental, engineered, and host-associated ecosystems where endospores constitute one of the most abundant cell types on Earth^55^.

## RESULTS

### Distribution of sigma factors in phage genomes

We characterized the diversity and distribution of sigma factors in more than 3,400 phage genomes in the viral orthologous groups (VOG) database. From this, we found that 14% of all the genomes analyzed contained at least one sigma factor gene while some genomes (0.7%) contained up to three different homologs (Fig. 1). The distribution of phage-encoded sigma factors was not random with respect to virus or host taxonomy (Fisher’s Exact Test, *P_simulated_* < 0.0001, Fig. 1). Sigma factor containing phages were most prevalent among those that infect the *Cyanobacteria* (65%), *Proteobacteria* (15%) and *Firmicutes* (19%). Among the tailed phages (*Caudovirales*), sigma factors were recovered in 5 of 9 families. Three of these five viral families (*Siphoviridae*, *Podoviridae*, and *Myoviridae*) have members that infect hosts from multiple bacterial phyla. However, only among phages that infect *Firmicutes* do all three of these virus families have members with sigma factor genes (Fig. 1c).

**Fig 1.**
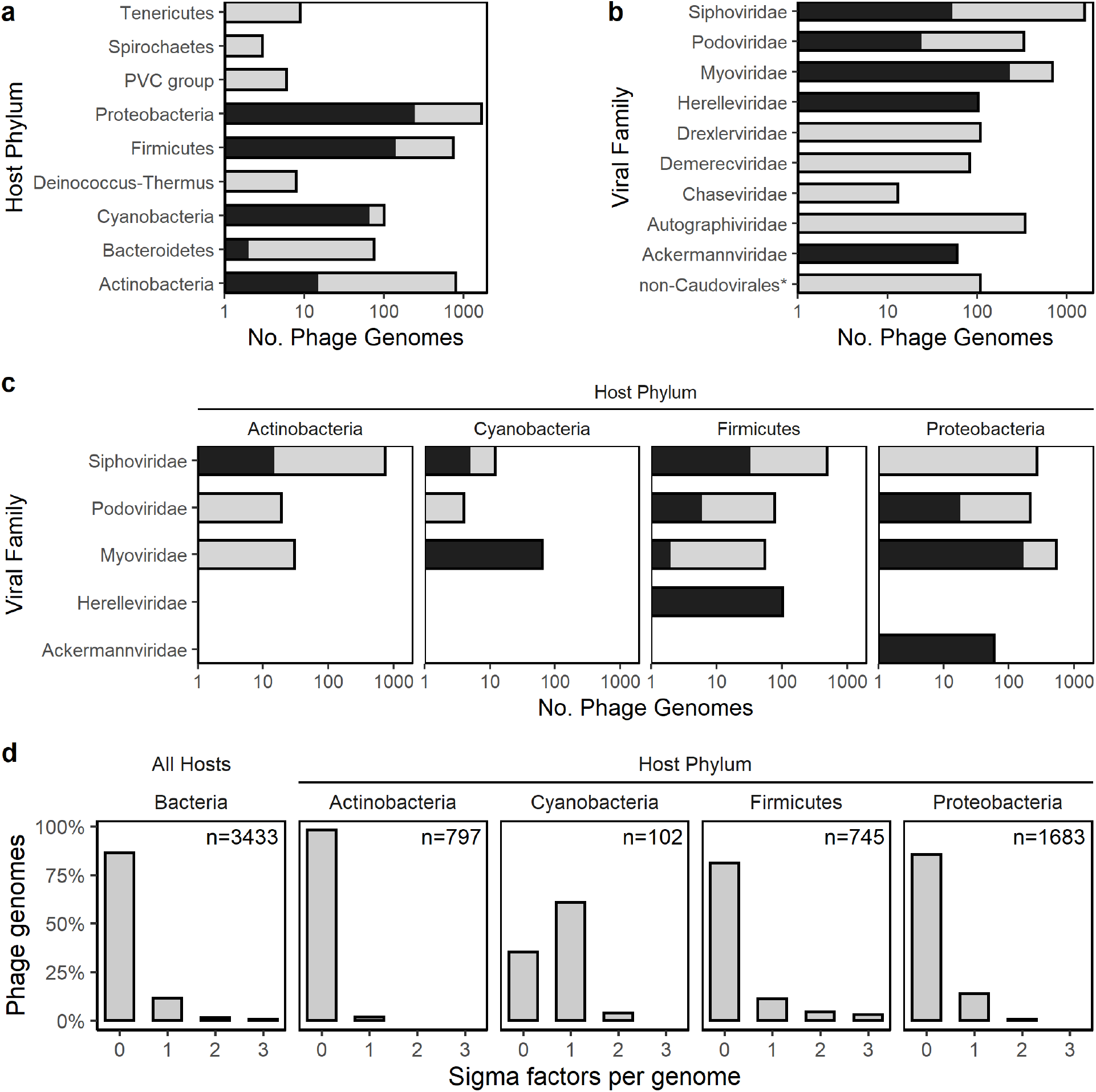
Sigma factors in viral genomes. **a-c**, The overall number of phage genomes in each taxonomic group is depicted by light grey bars with nested darker bars depicting how many of those genomes contained one or more sigma factors. Taxonomic groups are host phylum (**a**), viral family (**b**), and viral families containing sigma factors separated by host phyla (**c**). **d**, Multiplicity of sigma factors genes in phage genomes. The distribution of sigma factor gene counts in phage genomes is shown for all phages in the Viral Orthologous Groups (VOG) database (left facet), and for each of the host phyla as in **c**. Phage genomes from the VOG database were mapped to hosts using the Virus-Host Database. Viral families are specified for tailed phages (*Caudovirales*), and all non-tailed phages are shown in a single category of non-*Caudovirales*, with an asterisk to indicate that this is not a true viral family. The two *Bacteroidetes* phages with sigma factors indicated in **a**, each from a different viral family are excluded from **c** for clarity.

*Firmicutes* phages were also notable in that many of them encode multiple sigma factors. For phages that infect *Cyanobacteria* and *Proteobacteria*, 95% of the genomes with a sigma factor only contained a single copy of such a gene (Fig. 1d). In contrast, for phages that infect *Firmicutes*, 41% of the genomes with a sigma factor possessed two or three gene copies. Sigma factor gene multiplicity was detected in diverse *Firmicutes* phages, predominantly among strains that could infect spore-forming genera (Fig. S1). Most of these phages belonged to the *Herelleviridae*, a family of strictly lytic phages that was recently split from the *Myoviridae* and are thought to only infect bacteria in the *Firmicutes*^56^. Of the 102 *Herelleviridae* phages in our dataset, half possessed multiple sigma factors (n = 51) and only infected *Bacillus* hosts (Fig. S2). In contrast, none of the 49 phages that infect non spore-forming genera (*Enterococcus*, *Lactiplantibacillus*, *Lactobacillus*, *Listeria*, *Staphylococcus*) possessed more than a single sigma factor gene. Additionally, a similar trend was found among *Firmicutes* phages belonging to the *Siphoviridae*. Within this group, there were five phages with multiple sigma factors (Fig. S2), four that infect hosts capable of forming endospores (*Bacillus* and *Brevibacillus*), and one whose host range likely evolved from a *Bacillus* to *Staphylococcus* host^57^.

### Sporulation-specific sigma factors are encoded by *Firmicutes* phages

Many of the sigma factors recovered in *Firmicutes* phages were similar to the bacterial-encoded sigma factors involved in sporulation. To interpret this finding in an evolutionary context, we constructed a phylogeny of the VOG sigma factors together with sigma factors from diverse bacteria. Sporulation-specific sigma factors from bacteria belonged to a monophyletic clade that also contained *sigB*, a gene that *B. subtilis* uses to regulate its stress response (Fig. 2a,b). While some phage-encoded sigma factors were found in phage-specific clades, others clustered with bacterial-encoded sigma factors, including those that are known to regulate sporulation. All phage-encoded sigma factors in the sporulation clade were from *Firmicutes* phages, apart from two cyanophage genes that grouped with the *sigB* sub-clade.

**Fig 2.**
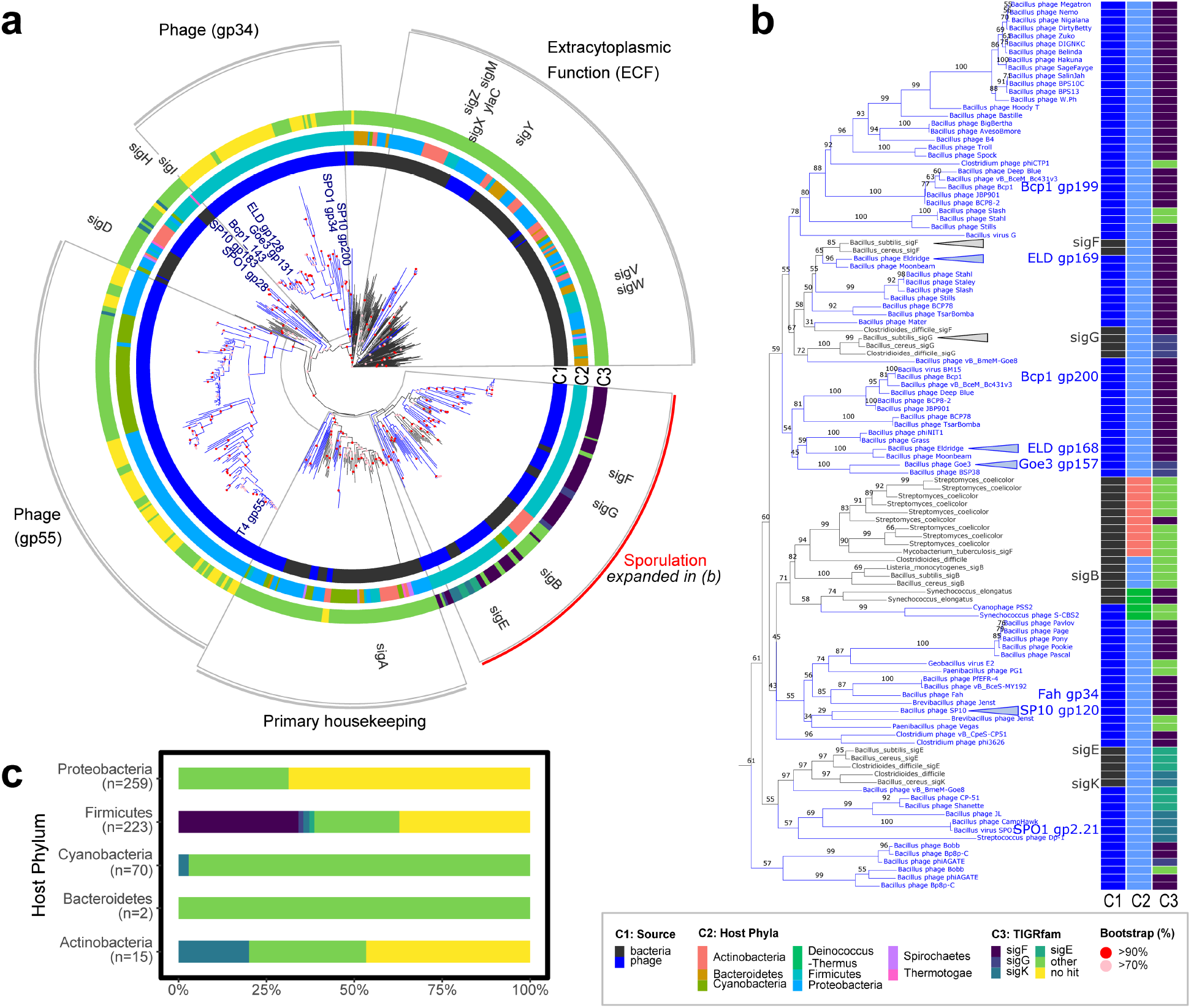
Phylogeny and classification of phage-encoded sigma factors. **a**, Phylogenetic tree representing proteins of phage-encoded sigma factor from the database of viral orthologous groups. For reference, we included sigma factors from 24 genomes of diverse bacterial species. Blue branches on the tree correspond with monophyletic clades that only contain phage-encoded sigma factors. Black branches show bacterial-encoded sigma factors and the internal branches leading to them. Phage proteins discussed in the text are labeled at branch tips (e.g., ELD gp128). In circle 1 (C1) we identify the source (phage vs. bacteria) of a sigma factor protein. In circle 2 (C2), we designate the taxonomy of the bacterial host (phyla). In circle 3 (C3) we provide the best hit (hmmscan sequence E-value) of the homolog to bacterial-encoded sigma factors families in TIGRfam. Outside of circle 3, we indicate the relative positions of *Bacillus subtilis* sigma factor genes and depict the different sigma factor families (grey wedges). Branch nodes are labeled with nonparametric bootstrap support values (n = 500). **b**, Clade of sporulation-specific sigma factors depicted by red arc in **a**. Triangles point to genes cloned and expressed in this study. **c**, Summarized proportions of phage-encoded sigma factors as a function of host taxonomy and protein family as described for circles 2 and 3 of panel **a**, respectively.

To complement the phylogenetic analyses, we classified phage-encoded sigma factors by estimating homology to bacterial sigma factor protein families (TIGRFAMs) using hidden Markov models. Phage-encoded homologs of sporulation-specific sigma factors of *Firmicutes*, henceforth “sporulation-like” sigma factors, were found nearly exclusively in phages that infect hosts of this phylum (Fig. 2c). Homologs of three of the four sporulation-specific sigma factors (*sigF*, *sigG* and *sigE*) were only detected in *Firmicutes* phages, with the majority resembling *sigF*. Furthermore, these homologs were only recovered from phages that infect *Bacillus*, *Clostridium*, and *Brevibacillus*, which are genera that commonly engage in endosporulation (Fig. S4). However, we did identify homologs of the sporulation-specific *sigK* in a few phages that infect *Cyanobacteria* and *Actinobacteria*.

The sporulation-like sigma factors recovered from phage genomes were distinct from phage-encoded sigma factors that regulate expression during lytic replication (Fig. 2a). In our phylogenetic analysis, sigma factors that are essential for phage development clustered in one of three other groups. First, there was a phage-specific clade of genes from phages that infect *Proteobacteria* and *Cyanobacteria*. This clade included gp55 from phage T4, which is known to control transcription during late infection stages^50,53^. Second, we identified a phage-specific clade of genes from *Firmicutes* phages, which included gp34, a sigma factor that controls late gene expression in *B. subtilis* phage SPO1^36^. Last, there was a paraphyletic cluster that included phage- and bacterial-derived sigma factors. This group contained gp28, which regulates the expression of middle infection genes in phage SPO1^36^. In addition, the latter group contained alternative sigma factors that are involved in bacterial regulation of motility (*sigD*), stationary phase (*sigH*), and heat stress response (*sigI*).

The pattern of gene multiplicity among *Firmicutes* phages (Fig. S1) reflects that these viruses contain sigma factors with divergent functions. They either cluster with genes known to regulate the lytic cycle or with bacterial sporulation-specific sigma factors (Fig. 2b). For example, *B. subtilis* phage SPO1 encodes two sigma factors that are essential for phage replication (gp28 and gp34)^36^. Likewise, phage SP10 has two sigma factors that cluster with the essential SPO1 genes which are known to regulate phage genes^58^. Each of these phages also encode a third homolog that clusters with sporulation-specific sigma factors. Similarly, all but two of the 58 *Firmicutes* phages exhibiting gene multiplicity contain one or two sporulation-like sigma factors (Fig. S3). Additionally, in several *Bacillus* phages the sole sigma factor is sporulation-like. This group of phages was enriched with representatives from the *Podoviridae* and the *Siphoviridae*, which includes some temperate phages such as Wβ which infects *B. anthracis*^42^. In sum, our bioinformatic analyses revealed that homologs of sporulation-specific sigma factors are found in diverse phages with contrasting lifestyles (i.e., lytic and temperate), often alongside sigma factors that phages use to regulate lytic reproduction.

### A sporulation-like sigma factor is non-essential for phage reproduction

Over time, a host-derived sigma factor could be repurposed by phages for regulating phage genes that are essential for reproductive functions like genome replication and capsid assembly. To test this hypothesis, we used CRISPR-Cas9 to delete the sporulation-like sigma factor g120 from SP10, a phage that infects *B. subtills* (see Figs. 2b and Table S1) and has overall 3 sigma factors (see above). After removing the entire coding sequence of the sporulation-like sigma factor g120, this phage could still productively infect its host, demonstrating that g120 was non-essential for phage reproduction under standard lab conditions. In fact, there was no detectable reduction of virulence when infecting *B. subtilis* with the mutant phage (t_12.9_ = 1.22, *P* = 0.25; Fig. S4).

### Phage-encoded sigma factors alter expression of host sporulation genes

To evaluate the ability of phage-encoded homologs to activate transcription of host sporulation genes, we cloned four diverse sporulation-like sigma factors (Fig. S5) from three *Bacillus* phages (Table S1) under an IPTG inducible promoter into the chromosome of a spore-forming strain of *B. subtilis*. As a control, we independently cloned the host-encoded sigma factors *sigF* and *sigG* into the same strain of *B. subtilis* in a similar manner. We then induced the expression of each of the cloned sigma factors during exponential growth, a time when native sporulation-specific sigma factors (i.e., *sigF*, *sigG, sigE* and *sigK*) and other sporulation genes are not typically expressed (Fig. 3a). Using RNAseq, we found that there was a strong positive correlation (ρ = 0.76) between differential gene expression following induction of the sigma factor g169 derived from phage Eldridge (ELDg169) and host-derived sigma factors (*sigF* and *sigG*) suggesting conservation of gene function (Fig. 3b, S6). Similarly, when ELDg169 was induced, we observed the upregulation of genes involved in sporulation (*P* <0.0001; Figs. 3c, S7). However, not all phage-derived sigma factors affected host expression equally. For example, induction of the other sporulation-like sigma factor cloned from phage Eldridge (ELDg168) also resulted in differential expression of many host genes. However, the genes affected by ELDg168 were significantly different than those observed in populations where host-derived sigma factors were induced (*P* < 0.0001; Fig. S6), and they were not enriched in sporulation genes (Figs. 3c, S7). Meanwhile, induction of the two other less conserved sporulation-like sigma factors (SP10 g120 and Goe3 g157) had only modest effects on gene expression, with less than 50 genes differentially expressed by each induced gene, and no enrichment of sporulation genes (Figs. 3c, S6, S7, S8).

**Fig 3.**
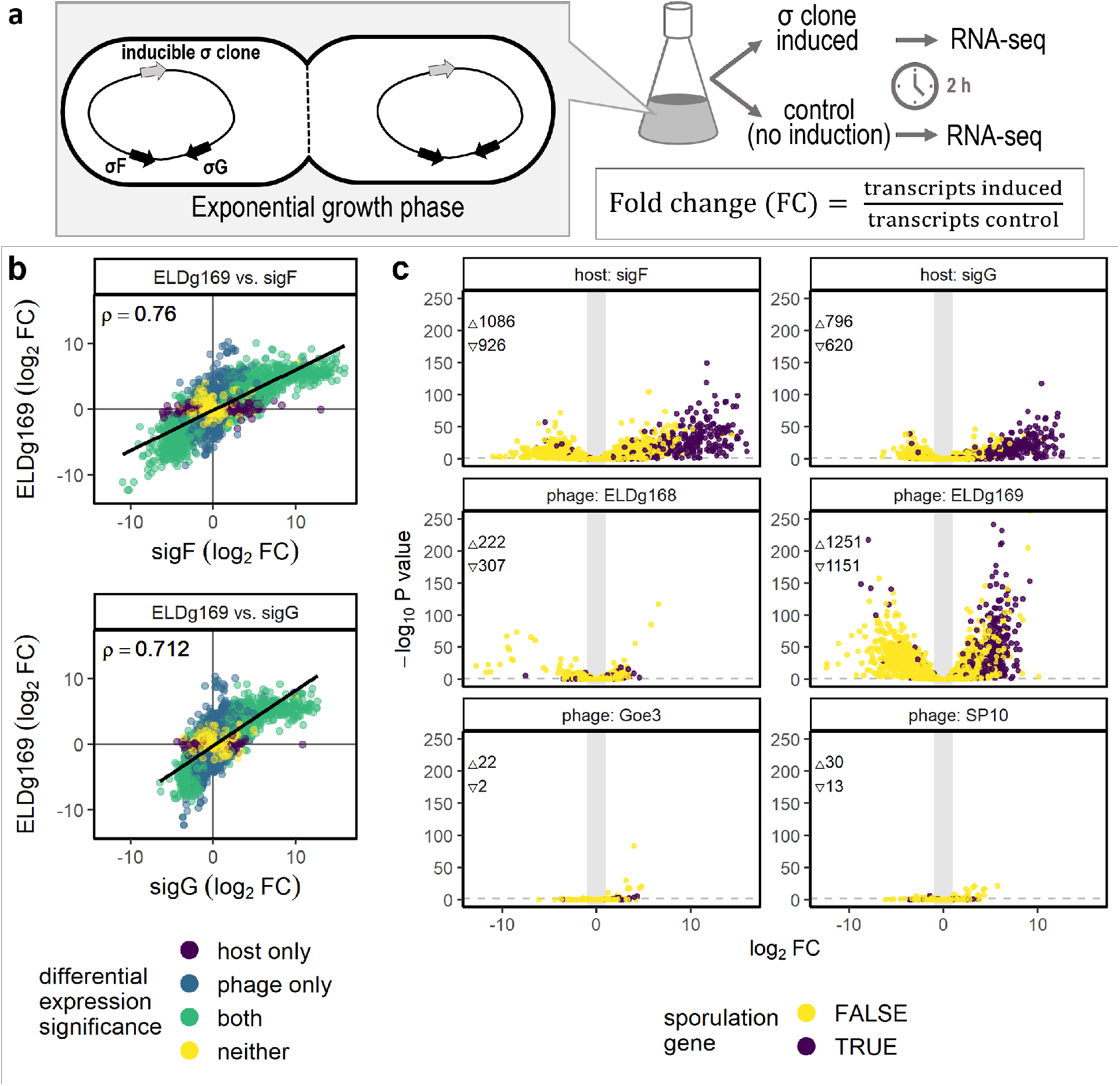
Bacterial gene expression following induction of phage-encoded sigma factors. **a**, Illustration of the experimental design. Phage and host derived sigma factors were cloned in *Bacillus subtilis* under an IPTG-inducible promoter (grey arrow) in a strain (Δ6) that contains a fully functional sporulation gene network, including its native sporulation specific sigma factors (black arrows show representative examples). To estimate differential expression, we compared splits of *Bacillus subtilis* cultures. One half was induced to express a cloned sigma factor during exponential growth, while the other half used as a non-induced control (n = 3). Differentially expressed genes were defined as those where *P* values < 0.05 (horizonal dashed line in **b**) and |fold change| > 2 (outside of vertical grey bar in **b**). **b**, Correlation of differentially expressed genes for cells induced to express phage-encoded sigma factors and cells induced to express bacteria-encoded sigma factors (*sigF* and *sigG*). Spearman’s correlation coefficient (*ρ*) is displayed. **c**, Sporulation genes^86^ were upregulated after induction of a phage-encoded sigma factor (ELDg169) and host-encoded sigma factors that regulate sporulation (*sigF* and *sigG*). The number of significantly upregulated (∆) and downregulated (∇) genes are noted.

### Phage-encoded sigma factors inhibit sporulation

Induction of the sporulation-like sigma factors altered spore yield in populations of *B. subtills* (Figs. 4, S9). When the host-derived sigma factors were induced in sporulating cultures, we observed a reduction in spore yield, a pattern that likely resulted from the misregulation of the sporulation gene network. Compared to an empty vector control with no cloned sigma factor, induction of *sigF* led to an 85% reduction in spore yield (t_8.2_ = 6.43, *P* < 0.001) while *sigG* reduced spore yield by 50% (t_13.9_ = 6.43, *P* = 0.015). Expression of phage-derived sigma factors also affected sporulation, in one case to a greater degree than host-derived sigma factors (Fig. 4, Table S2). Induction of ELDg169 reduced the spore yield by 99% compared to the empty vector control (t_7_ = 7.8, *P* < 0.0001), while the other Eldridge-derived gene, ELDg168, had a smaller (∼33%) and marginal effect on spore yield (t_15.1_ = 2.1, *P* = 0.064). The sigma factor from phage Goe3 reduced spore yield by > 50% (t_8.9_ = 4.39, *P* = 0.002), while expression of the sigma factor from phage SP10 had no effect on spore yield (t_10.4_ = 0.46, *P* = 0.65). Because spore yield was calculated as a percentage of the total population, we compared cell counts between induced and non-induced controls (Fig. S10). From this, we concluded that the observed reductions in spore yield were not due to a significant reduction in vegetative cells (Table S3).

**Fig. 4.**
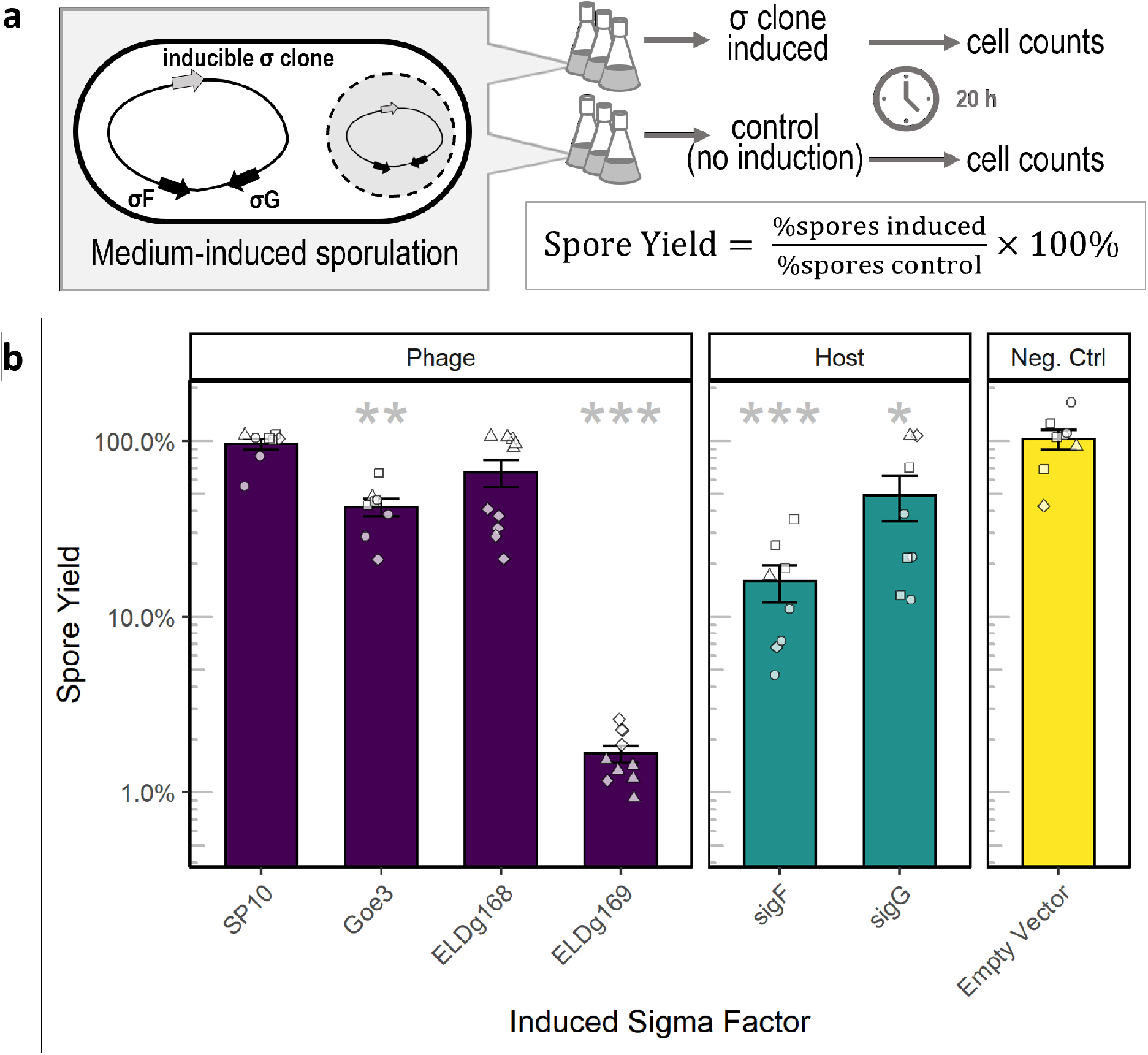
Phage-derived sigma factors reduced sporulation. **a**, Illustration of the experimental design. Phage and host derived sigma factors were cloned in *Bacillus subtilis* under an IPTG-inducible promoter (grey arrow) in a strain (Δ6) that contains a fully functional sporulation gene network, including its native sporulation specific sigma factors (black arrows show representative examples). We compared sporulation in these strains to an empty-vector negative control strain that also had an IPTG-inducible promoter. For each data point in **b** replicate cultures (n = 6) of a single colony were grown in sporulation medium for 4.5 hours, a time at which sporulation was induced by nutrient exhaustion. We then induced expression of the cloned sigma factor by the addition of IPTG to half of the replicates, while leaving the remainder as controls. Spore and vegetative cells were quantified by flow-cytometry. Spore yield was calculated as the ratio of percent spores in induced cultures and their paired controls. **b**, Bars represent the mean ± SEM of independent clones (n ≥ 8). Shapes represent different experimental batches. Asterisks correspond to significance levels (adjusted for multiple testing) from Welch’s *t*-tests used to evaluate the effect of each sigma factor compared to the empty-vector negative control (* = *P* < 0.05, ** = *P* < 0.01, *** = *P* < 0.001). ELD = phage Eldridge.

## DISCUSSION

It is well known that some phages encode for sigma factors that regulate transcription during lytic development^50,52,53^. In fact, early investigations of phage-encoded sigma factors elucidated how the swapping of sigma subunits controls gene transcription by RNA polymerase^50^. However, those phage-encoded sigma factors are quite divergent from bacteria-encoded sigma factors^53,59^. More recently, genomic data have identified phage-encoded sigma factors that bear a greater resemblance to bacterial sigma factors, especially those involved in the regulation of sporulation in the *Firmicutes*. While the function of these homologs remains unexplored, it is possible that sporulation-like sigma factors have been coopted by phages to manipulate host dormancy in ways that could potentially enhance their reproduction and survival. In support of this hypothesis, our bioinformatic and phylogenetic analyses revealed that sporulation-like sigma factors were found primarily in phages that infect spore-forming hosts, where they could potentially act as analogs of the host’s own sporulation-specific sigma factors. We show that these phage-encoded sigma factors are distinct from genes known to be essential for regulating lytic reproduction and that they are non-essential for phage reproduction. When expressed in *Bacillus subtilis,* phage-encoded sigma factors reduced spore yield, likely owing to their ability to alter the expression of host genes, which in one case led to transcriptional activation of the sporulation network. Together our findings highlight novel ways in which dormancy may influence antagonistic coevolution between spore-forming bacteria and their phages.

### Diversity and distribution of phage-encoded sigma factors

Phages are able to regulate gene expression in different ways. For example, they can make use of host-encoded sigma factors, encode sigma factor analogs (e.g., gp33 in T4), or use their own RNA polymerases that do not require sigma factors^50,53,60,61^. In addition, some phages encode their own sigma factors. Of the nearly 3,500 genomes examined, we found that 14% of the phages possess a sigma factor. The distribution of sigma factors in phage genomes was strongly influenced by host and viral taxonomy. While common among phages that infect *Cyanobacteria* and *Proteobacteria*, only among the *Firmicutes* phages did we recover sigma factors that were encoded by representatives from all three of the major tailed-phage families (*Siphoviridae*, *Podoviridae* and *Myoviridae*; Fig. 1). This suggests that sigma factors may be beneficial to diverse phages that infect *Firmicutes* hosts. Furthermore, we identified a host-specific pattern of gene multiplicity. A diverse set of *Firmicutes* phages carried two sigma factors with host ranges that were limited to spore-forming genera, while phages containing three sigma factors were restricted to just two viral families (*Herelleviridae* and *Siphoviridae*) that could only infect strains of *Bacillus*. This convergent pattern of gene multiplicity is associated with the presence of sporulation-like sigma factors in phages that infect *Firmicutes*. Given the high degree of sequence similarity between sigma factors encoded by phages and those encoded by their hosts, it seems likely that viruses have acquired these genes from their hosts. Furthermore, because the genes are widespread among diverse viral lineages, it seems reasonable to hypothesize that there are functional consequences for phage that encode sporulation-like sigma factors. Nevertheless, in the database used in our study, these genes are not universal in phages that infect spore-forming *Firmicutes*. We expect that future investigations of metagenomes from environmental, engineered, and host-associated ecosystems will shed light on the eco-evolutionary factors that influence the distribution and abundance of dormancy related auxiliary genes, including sporulation-like sigma factors.

### Sporulation-like sigma factors appear to be non-essential for phages

In bacteria, sporulation and other specialized functions are controlled by alternative sigma factors that replace the primary housekeeping sigma factor^51^. Because alternative sigma factors are only transiently used, their deletion is typically non-lethal and thus they are considered non-essential. Similarly, multiple lines of evidence suggest that sporulation-like sigma factors are non-essential for fundamental aspects of phage biology. Unlike gp28 and gp34, which are required by phage SPO1, a sporulation-like sigma factor (gp2.21) is known to be non-essential for lytic infection under standard laboratory conditions^36^. While it has been hypothesized that gp2.21 directs transcription of phage genes from *sigK*-like promoters found in its genome^36^, this has not been confirmed. In our study, we deleted the sporulation-like sigma factor from phage SP10 (g120) with no observable effect on virulence (Fig. S4). Even though g120 is expressed during SP10 infection of its host^58^, available evidence suggests that at least some sporulation-like sigma factors are non-essential for lytic infection. If this pattern is generalizable, then non-essentiality could reflect that phage-encoded sigma factors have alternate function, which could involve the control of genes that are required for processes other than replication.

### Phage-encoded sigma factors disrupt regulation of the sporulation network

The prevalence of sporulation-like sigma factors among diverse phages suggests that these genes may have consequences for phage performance. One possibility is that these sigma factors play the same role in phages as they do in their host, that is, they regulate the expression of sporulation genes. In support of this notion, previous experimental studies have documented that phage-encoded sporulation-like sigma factors retain a function that is relevant to host sporulation. For example, in lysogenic *B. anthracis*, expression of a sporulation-like sigma factor encoded by the Wβ prophage is elevated during sporulation^42^. Additionally, *in vitro* reconstitution of RNA polymerase with the sporulation-like sigma factor of *B. anthracis* phage Fah, a close relative of Wβ, revealed patterns of transcriptional activity and inhibition that are similar to that of a bacterial-encoded sigma factor (*sigF*)^54^. Finally, sporulation-like sigma factors cloned from two diverse *B. anthracis* phages, Bcp1 (*Herelleviridae*) and Wip4 (*Siphoviridae*), were associated with phage-dependent inhibition of host sporulation^41^.

In our study, the induction of phage-derived sporulation-like sigma factors altered host gene expression and disrupted sporulation in *B. subtilis*. The magnitude of this effect was contingent upon the identity and phylogenetic distance between the phage- and host-encoded sigma factors (Fig, S11). When we induced ELDg169, the most host-like gene in our experiments, we observed the differential expression of hundreds of bacterial genes, including the upregulation of nearly 400 sporulation genes (Fig. S7), which led to a ∼99% reduction in spore yield (Fig. 4b). In contrast, we observed a less pronounced reduction in spore yield, and almost no effect on the sporulation gene network when two less conserved phage-derived genes (ELDg168 and Goe3 g157) were induced. Last, induction of g120 from phage SP10 resulted in a very mild transcriptional response and had no effect on spore yield, in accordance with it being the most divergent gene from bacterial *sigF* and *sigG*. Thus, our studies suggest that that the functionality of phage-encoded sigma factors is influenced by the degree of similarity with host-encoded sigma factors. However, this does not rule out that phage-encoded sigma factors may play other roles depending on host and environmental conditions. For example, the manipulation of sporulation seems to be the likely function of these genes in phage Eldridge, possibly also in phage Goe3, but not in phage SP10.

As is the case with many evolutionary investigations, carefully planned experiments are required to make strong inferences about the adaptive significance of genetic variation in a population. We hypothesize that sporulation-like sigma factors could influence phage fitness in different ways. By controlling the timing of host sporulation, phages may increase their probability of entrapment^46^, which would increase the survivorship component of phage fitness. On the other hand, by impeding sporulation, there is more opportunity for virus replication, which would increase the reproductive component of phage fitness. In our control experiments, induced expression of bacterial genes activated the sporulation network during exponential growth. However, when induced during sporulation, these host genes, which are known to promote sporulation, led to a reduction in spore yield. This pattern likely stems from the regulatory genes being expressed in the wrong place and time. As a consequence, it is challenging to conclusively deduce whether the reduction in spore-yield reflects the function of phage proteins in infected cells or other aspects of our experiment. However, results from one particular phage provide some additional insight. The two Eldridge-derived genes, which are genomic neighbors, both reduced sporulation to some degree, even though they had very different transcriptional profiles. If these adjacent genes work toward a common function, it seems likely that this results in the inhibition of host sporulation. This interpretation is consistent with the observation that complete inhibition of sporulation in *B. anthracis* is likely mediated by a pair of sporulation-like sigma factors found in tandem in the genome of phage Bcp1^41^. Additional experiments are needed to understand the evolutionary consequences of sporulation-like sigma factors for phage populations. For example, comparisons between phages in which sigma factors are present or deleted (like the SP10 mutant we constructed) will be useful in determining whether auxiliary dormancy genes enhance phage fitness. Likewise, comparative transcriptomic analysis during infection with phages with and without sigma factors may help identify the regulatory targets of phage-encoded sigma factors. Looking beyond sporulation, studies are needed to better understand how bacterial-like sigma factors may be used by phages to manipulate other survival strategies that are common in non-growing bacteria^62^.

Taken together, our findings reveal a pattern of genomic convergence. Phages from diverse families, with contrasting infection strategies (virulent vs. temperate) that infect spore-forming hosts also contain sporulation-specific sigma factor homologs. Importantly, these sigma factors are phylogenetically distinct from sigma factors that regulate phage genes during lytic replication. At least some of these genes are non-essential for phages, which further supports the view that alternate sigma factors were not acquired for the purpose of regulating lytic programs. Instead, our experiments demonstrate that phage-encoded sigma factors retain features of their ancestral function, which means that they can be viewed as auxiliary genes that can regulate the sporulation network with consequences for spore yield. That being said, our results also reveal that some sporulation-like sigma factors in phages have variable and yet unknown functions, which may reflect neutral or adaptive divergence.

### Eco-evolutionary implications of auxiliary dormancy genes

As obligate parasites, phages are unavoidably dependent on the metabolism of their bacterial hosts. Bacteria are capable of responding to their dynamic environments by replacing the sigma subunit of RNA polymerase, which leads to changes in gene expression^50,51^. Some phages use a similar strategy to coordinate expression of their own genes during different stages of infection^53^. Our analysis points to the existence of a second class of sigma factors that phages may use to manipulate host metabolism. Whether phages promote or inhibit sporulation, such manipulation of host dormancy has the potential to modify the environmental conditions under which we should expect to find bacteria in an active vs. dormant state. This has implications for development of novel therapeutic treatments that combine phage therapy with antimicrobials, which tend to target metabolically active bacteria^63^.

Viruses have evolved to utilize, maintain, and rearrange a variety of biochemical pathways in the cells that they take over^25,28,64^. The discovery of auxiliary metabolic genes has revealed that viruses can appropriate the cellular building blocks and protein translation machinery of their hosts. Our findings highlight an additional aspect of phage-host coevolution involving the co-option of gene networks used to coordinate a complex and ancient form of dormancy, which many microorganisms use to contend with harsh and unpredictable environments. Manipulation of the host’s response to such conditions through the acquisition of host regulatory genes could represent a strategy which buffers viruses from the dynamic cellular environment on which their survival and reproduction is dependent.

## METHODS

### Phage sigma factor distribution and classification

We retrieved sigma factors from the database of viral orthologous groups (VOG release vog202, vogdb.org) based on text searches of the VOG descriptors (Table S4). We matched VOG proteins to host and viral taxonomy using the virus-host database^65^. We classified phage-encoded sigma factors using hmmscan with default parameter settings using HMMER v3.3^66^, and queried each protein against hidden Markov model (HMM) profiles of bacterial-encoded sigma factor families that were retrieved from TIGRFAM (Table S5). We used the best hmmscan match (smallest sequence E-value) to classify proteins, unless it was a general TIGRFAM (“sigma70-ECF” or “SigBFG”), in which case the next best match was chosen, if available.

### Phylogenetic analysis

For phylogenetic analysis of sigma factors, we aligned VOG proteins from phage genomes along with sigma factor proteins belonging to 24 bacteria from diverse taxa^67^. We aligned sigma factor sequences using MAFFT (v.7.475)^68^ with the E-INS-I strategy and trimmed the alignment with trimAL (v1.4.rev22)^69^ using the gappyout method. From the 180 amino acids in the trimmed alignment we then inferred 200 maximum likelihood phylogenetic trees using RAxML-NG (v0.9.0-pthreads)^70^ with the LG+G4 substitution model selected using modeltest-NG (v0.1.6)^71^ with default settings. We present the best scoring maximum likelihood tree with Transfer Bootstrap Expectation supports^72^ from 500 bootstrapped trees. We plotted the tree using the ggtreeExtra R package^73^.

### Strains and media

Strains used are listed in Table S1. For routine culturing of bacteria, we used LB medium with low salt (5 g/L NaCl). We amended this recipe with agar (15 g/L) for plating and with CaCl_2_ (10 mM) to facilitate virus adsorption. We used Difco sporulation media (DSM) for sporulation assays^74^. For plaque assays, we used double-layer plating with 0.3% agar overlays^75^. To amplify phages, we collected lysates from plate infections after flooding Petri dishes with phage buffer (10 mM Tris, 10 mM MgSO_4_, 4 g/L NaCl, 1 mM CaCl_2_, pH 7.5). We then cleared the phage-containing buffer from bacteria by centrifugation (7,200 Xg, 10 min) and filtration (0.2 μm).

### Deletion of phage-encoded sigma factor

We used the CRISPR-Cas9 system and the CutSPR assay design-tool^76^ to test whether sigma factors are essential for phage replication. Briefly, we cloned a single-guide RNA and a deletion cassette into plasmid pJOE8999^77^ (Tables S6, S7) and transformed the resulting plasmid into *B. subtilis* TS01 (Table S3), which was made competent with D-mannitol induction. We next infected the transformed culture with phage SP10 (Table S1) and conducted a plaque assay with medium containing the Cas9-inducer D-mannose. Using primers SP10_validF+R (Table S7), we screened multiple plaques for the deletion. To isolate the mutant phages, we picked and replated PCR-positive plaques onto host *B. subtilis* Δ6^78^ (Table S1). We screened these secondary plaques as above and confirmed the deletion by Sanger sequencing of the locus. We then quantified the virulence of the mutant and wild-type phages^79^. After dispensing *B. subtilis* Δ6 host cultures (OD600 = 1) into microtiter wells, we infected cells with serially diluted lysates of SP10 or SP10 Δg120 that were adjusted to an equal titer. We monitored bacterial density during growth for 16 h by OD600 with a Synergy H1 plate reader (Biotek). From this, we calculated the virulence index^79^ based on change in bacterial growth and lysis as a function of the phage:bacteria ratio (i.e., multiplicity of infection; Fig S4).

### Inducible expression of sigma factors

We tested the effect of phage-derived sigma factors on bacterial expression by cloning coding sequences under an inducible promoter into an ectopic site (*amyE*) of the *B. subtilis* genome. As a control, we also cloned host-derived sporulation genes (s*igF* and *sigG*) in the same manner, and a gene-less promoter as a negative control.

#### Strain construction

We amplified coding sequences by PCR from phage lysates or from extracted bacterial genomic DNA as templates using primers adapted with restriction sites, and a ribosome binding site on the forward primer (Table S7). We then cloned the PCR products into plasmid pDR110 (Table S6) by restriction enzyme digestion (Table S6), gel purification, and ligation (T4 ligase). We selected for plasmids that were transformed into *E. coli* (One Shot TOP10, Fisher) with ampicillin (100 µg/ml) and verified the insertion by PCR and Sanger sequencing using primers oDAS9+10 (Table S7). We transformed purified plasmids (QIAgen mini prep) into *B. subtilis* TS01, as described above, using spectinomycin selection (100 μg/ml). We verified the insertion into the *amyE* locus by PCR and Sanger sequencing, and by the loss of erythromycin resistance carried in the *amyE* locus by strain TS01.

#### Transcriptional response to phage-encoded sigma factor

We diluted overnight *B. subtilis* cultures (OD600 = 0.1) in fresh LB and grew them (37 °C, 200 RPM) to mid exponential phase (OD600 = 0.5). We then split the cultures and added 1mM IPTG (final concentration) to one half to induce expression of the cloned gene. We added an equal volume of water to the other half of the split culture as a non-induced control. After induction, we incubated the cultures for 2 h before harvesting cells. Upon sampling, we immediately treated bacteria with the RNAprotect Bacteria reagent (Qiagen), and stored pellets at −80 °C for <1 week before RNA extraction using RNeasy Protect Bacteria Mini Kit (Qiagen) according to the manufacturer’s instructions (protocol #5), including an on-column RNase-free DNase digestion. Library construction, sequencing, and analysis of differential gene expression were all carried out at the Indiana University Center for Genomics and Bioinformatics. Libraries were constructed using the Illumina TruSeq Stranded mRNA HT kit following depletion of rRNA using Illumina Ribo-Zero Plus kit. Libraries were then sequenced on an Illumina NextSeq 500 platform as paired end reads (2 x 38 bp). We trimmed adapters and filtered reads using Trimmomatic 0.38^80^ with the cutoff threshold for average base quality score set at 20 over a window of 3 bases. Reads shorter than 20 bases post-trimming were excluded. We mapped the cleaned reads to the reference genome (Deposited with sequencing data to the Gene Expression Omnibus, see below) using bowtie2 version 2.3.2^81^, and counted reads mapping concordantly and uniquely to the annotated genes using featureCounts tool ver. 2.0.0 of subread package^82^. Read alignments to antisense strand, or to multiple regions on the genome or those overlapping with multiple genes were ignored (parameters: -s 2 -p -B -C). We performed differential expression analysis using DESeq2 ver. 1.24.0^83^ from normalized read counts by comparing samples induced with IPTG to non-induced paired control samples, with multiple-testing correction. We tested for the effects of gene enrichment and overlap of differentially expressed genes using the hypergeometric distribution in R^84^.

#### Sporulation of cells expressing cloned sigma factors

To test for the effects of induced sigma factors on host sporulation, we diluted overnight *B. subtilis* cultures in fresh DSM (OD600 = 0.05) and dispensed each culture into multiple wells of a 96-well plate that was then incubated in a Biotek Synergy H1 plate reader (37 °C, fast and continuous shake setting). Under these conditions, we determined that cells enter stationary phase after approximately 4.5 h, marking the onset of sporulation. At this time, we induced expression of the cloned gene by adding IPTG (final concentration 1 mM) to half the cultures in the plate. We added water to the rest of the wells, which served as non-induced controls. At 24 h, we quantified the number of spores and vegetative cells in each well using a flow-cytometry assay that distinguished spores from vegetative cells (non-spores) based on differential uptake of the nucleic acid stain SYBR green^85^. We diluted each sample in TE buffer (pH 8) and then fixed the cells in 0.5% glutaraldehyde for 15 min at 4 °C. We stained the fixed samples with SYBR green (20,000x dilution of commercial stock, Lonza) for 10 min at room temperature in the dark. We then enumerated cells using a volumetric NovoCyte 2000R flow cytometer (Acea; ex 488 nm, em 530/30 nm) and an automatic gating pipeline.

### Code and data availability

All code and data used in the analyses in this study are available at github.com/LennonLab/sigma-spore-phage and github.com/LennonLab/sigma-spore-phage-flow. RNA sequencing data are available at the Gene Expression Omnibus under accession number GSE187004. In addition, prior to publication, all data and code will be made available on Zenodo.

## ACKNOWLEDGEMENTS

We acknowledge R Hertel, L Temple, AC Hernandez, M Liu, X Wang, David Rudner, D Zeigler, and DB Kearns for suggestions and strains; E Long and C Chen for technical support; and J Bird for assistance with preliminary computation. An earlier version of the manuscript was improved based on critical feedback from FJ Fishman, C Karakoç, A Magalie, JG McMullen, RZ Moger-Reischer, EA Muller, PG Wall and JS Weitz. Research was supported by the National Science Foundation (DEB-1934554 JTL and DAS, DBI-2022049 JTL), US Army Research Office Grant (W911NF-14-1-0411 JTL), the National Aeronautics and Space Administration (80NSSC20K0618 JTL). This research was also supported in part by Lilly Endowment, Inc., through its support for the Indiana University Pervasive Technology Institute.

## REFERENCES

1. Lennon, J. T., den Hollander, F., Wilke-Berenguer, M. & Blath, J. Principles of seed banks and the emergence of complexity from dormancy. Nat. Commun. 12, 1–16 (2021).

2. Lennon, J. T. & Jones, S. E. Microbial seed banks: the ecological and evolutionary implications of dormancy. Nat. Rev. Microbiol. 9, 119–130 (2011).

3. Cáceres, C. E. Temporal variation, dormancy, and coexistence: a field test of the storage effect. Proc. Natl. Acad. Sci. USA 94, 9171–9175 (1997).

4. Klobutcher, L. A., Ragkousi, K. & Setlow, P. The *Bacillus subtilis* spore coat provides “eat resistance” during phagocytic predation by the protozoan *Tetrahymena thermophila*. Proc. Natl. Acad. Sci. USA 103, 165–170 (2006).

5. Verin, M. & Tellier, A. Host-parasite coevolution can promote the evolution of seed banking as a bet-hedging strategy. Evolution 72, 1362–1372 (2018).

6. Berleman, J. E. & Bauer, C. E. Characterization of cyst cell formation in the purple photosynthetic bacterium *Rhodospirillum centenum*. Microbiology 150, 383–390 (2004).

7. Driks, A. & Eichenberger, P. in The Bacterial Spore: From Molecules to Systems 179–200 (American Society of Microbiology, 2016).

8. Kaplan-Levy, R. N., Hadas, O., Summers, M. L., Rücker, J. & Sukenik, A. in Dormancy and resistance in harsh environments Topics in Current Genetics (eds Esther Lubzens, Joan Cerdà, & Melody Clark) 5–27 (Springer, Berlin, Heidelberg, 2010).

9. Burroughs, N. J., Marsh, P. & Wellington, E. M. H. Mathematical analysis of growth and interaction dynamics of streptomycetes and a bacteriophage in soil. Appl. Environ. Microbiol. 66, 3868–3877 (2000).

10. Dowding, J. Characterization of a bacteriophage virulent for *Streptomyces coelicolor* A3 (2). Microbiology 76, 163–176 (1973).

11. Singh, R. N. & Singh, P. K. Isolation of Cyanophages from India. Nature 216, 1020–1021 (1967).

12. Gabiatti, N. et al. Bacterial endospores as phage genome carriers and protective shells. Appl. Environ. Microbiol. 84, e01186 (2018).

13. Hadas, H., Einav, M., Fishov, I. & Zaritsky, A. Bacteriophage T4 development depends on the physiology of its host *Escherichia coli*. Microbiology 143 ( Pt 1), 179–185 (1997).

14. Middelboe, M. Bacterial growth rate and marine virus–host dynamics. Microb. Ecol. 40, 114–124 (2000).

15. Abedon, S. T. & Yin, J. in Bacteriophages: Methods and Protocols, Volume 1: Isolation, Characterization, and Interactions (eds Martha R. J. Clokie & Andrew M. Kropinski) 161–174 (Humana Press, 2009).

16. Bryan, D., El-Shibiny, A., Hobbs, Z., Porter, J. & Kutter, E. M. Bacteriophage T4 infection of stationary phase *E. coli*: life after log from a phage perspective. Front Microbiol 7 (2016).

17. Makarova, K. S., Anantharaman, V., Aravind, L. & Koonin, E. V. Live virus-free or die: coupling of antivirus immunity and programmed suicide or dormancy in prokaryotes. Biol Direct 7, 7–40 (2012).

18. Koonin, E. V. & Zhang, F. Coupling immunity and programmed cell suicide in prokaryotes: Life-or-death choices. Bioessays 39, e201600186 (2017).

19. Meeske, A. J., Nakandakari-Higa, S. & Marraffini, L. A. Cas13-induced cellular dormancy prevents the rise of CRISPR-resistant bacteriophage. Nature 570, 241–245 (2019).

20. Bautista, M. A., Zhang, C. & Whitaker, R. J. Virus-Induced dormancy in the archaeon *Sulfolobus islandicus*. mBio 6, e02565 (2015).

21. Lenski, R. E. in Adv. Microb. Ecol. Vol. 10 (ed K. C. Marshall) 1–44 (Springer, 1988).

22. Lopatina, A., Tal, N. & Sorek, R. Abortive infection: bacterial suicide as an antiviral immune strategy. Annu Rev Virol 7, 371–384 (2020).

23. van Houte, S., Buckling, A. & Westra, E. R. Evolutionary ecology of prokaryotic immune mechanisms. Microbiol. Mol. Biol. Rev. 80, 745–763 (2016).

24. Ofir, G. & Sorek, R. Contemporary phage biology: from classic models to new insights. Cell 172, 1260–1270 (2018).

25. Thompson, L. R. et al. Phage auxiliary metabolic genes and the redirection of cyanobacterial host carbon metabolism. Proc. Natl. Acad. Sci. USA 108, E757–E764 (2011).

26. Lindell, D. et al. Transfer of photosynthesis genes to and from *Prochlorococcus* viruses. Proc. Natl. Acad. Sci. USA 101, 11013–11018 (2004).

27. Mann, N. H., Cook, A., Millard, A., Bailey, S. & Clokie, M. Bacterial photosynthesis genes in a virus. Nature 424, 741–741 (2003).

28. Hargreaves, K. R., Kropinski, A. M. & Clokie, M. R. Bacteriophage behavioral ecology: How phages alter their bacterial host’s habits. Bacteriophage 4, e29866–e29866 (2014).

29. Silpe, J. E. & Bassler, B. L. Phage-encoded LuxR-type receptors responsive to host-produced bacterial quorum-sensing autoinducers. mBio 10, e00638–00619 (2019).

30. Mizuno, C. M. et al. Numerous cultivated and uncultivated viruses encode ribosomal proteins. Nat. Commun. 10, 752 (2019).

31. Seed, K. D., Lazinski, D. W., Calderwood, S. B. & Camilli, A. A bacteriophage encodes its own CRISPR/Cas adaptive response to evade host innate immunity. Nature 494, 489–491 (2013).

32. Murphy, J. et al. Methyltransferases acquired by lactococcal 936-type phage provide protection against restriction endonuclease activity. BMC Genomics 15, 1–11 (2014).

33. Dragoš, A. et al. Pervasive prophage recombination occurs during evolution of spore-forming Bacilli. ISME J 15, 1344–1358 (2021).

34. Reveille, A. M., Eldridge, K. A. & Temple, L. M. Complete genome sequence of *Bacillus megaterium* bacteriophage Eldridge. Genome Announc. 4, e0172815 (2016).

35. Ritz, M. P., Perl, A. L., Colquhoun, J. M., Chamakura, K. R. & Everett, G. F. K. Complete genome of *Bacillus subtilis* myophage CampHawk. Microbiol Resour Announc 1, e00984–00913 (2013).

36. Stewart, C. R. et al. The genome of *Bacillus subtilis* bacteriophage SPO1. J. Mol. Biol. 388, 48–70 (2009).

37. Van Goethem, M. W., Swenson, T. L., Trubl, G., Roux, S. & Northen, T. R. Characteristics of wetting-induced bacteriophage blooms in biological soil crust. mBio 10, e02287–02219 (2019).

38. Yuan, Y., Gao, M., Wu, D., Liu, P. & Wu, Y. Genome characteristics of a novel phage from *Bacillus thuringiensis* showing high similarity with phage from *Bacillus cereus*. PLoS One 7, e37557 (2012).

39. El-Arabi, T. F. et al. Genome sequence and analysis of a broad-host range lytic bacteriophage that infects the *Bacillus cereus* group. Virol J 10, 1–11 (2013).

40. Zimmer, M., Scherer, S. & Loessner, M. J. Genomic analysis of *Clostridium perfringens* bacteriophage φ3626, which integrates into guaA and possibly affects sporulation. J. Bacteriol. 184, 4359–4368 (2002).

41. Schuch, R. & Fischetti, V. A. The secret life of the anthrax agent Bacillus anthracis: bacteriophage-mediated ecological adaptations. PLoS One 4 (2009).

42. Schuch, R. & Fischetti, V. A. Detailed genomic analysis of the Wβ and γ phages infecting *Bacillus anthracis*: implications for evolution of environmental fitness and antibiotic resistance. J. Bacteriol. 188, 3037–3051 (2006).

43. Sonenshein, A. L. Trapping of unreplicated phage DNA into spores of *Bacillus subtilis* and its stabilization against damage by ^32^P decay. Virology 42, 488–495 (1970).

44. Sonenshein, A. L. Bacteriophages: How bacterial spores capture and protect phage DNA. Curr. Biol. 16, R14–R16 (2006).

45. Takahashi, I. Incorporation of bacteriophage genome by spores of *Bacillus subtilis*. J. Bacteriol. 87, 1499–1502 (1964).

46. Sonenshein, A. L. & Roscoe, D. H. The course of phage ∅e infection in sporulating cells of *Bacillus subtilis* strain 3610. Virology 39, 265–276 (1969).

47. Galperin, M. Y. et al. Genomic determinants of sporulation in *Bacilli* and *Clostridia*: towards the minimal set of sporulation-specific genes. Environ. Microbiol. 14, 2870–2890 (2012).

48. Meeske, A. J. et al. High-throughput genetic screens identify a large and diverse collection of new sporulation genes in *Bacillus subtilis*. PLoS Biol. 14, e1002341 (2016).

49. Ramos-Silva, P., Serrano, M. & Henriques, A. O. From root to tips: sporulation evolution and specialization in *Bacillus subtilis* and the intestinal pathogen *Clostridioides difficile*. Mol. Biol. Evol. 36, 2714–2736 (2019).

50. Helmann, J. D. Where to begin? Sigma factors and the selectivity of transcription initiation in bacteria. Mol. Microbiol. 112, 335–347 (2019).

51. Paget, M. S. Bacterial sigma factors and anti-sigma factors: structure, function and distribution. Biomolecules 5, 1245–1265 (2015).

52. Losick, R. & Pero, J. Cascades of sigma factors. Cell 25, 582–584 (1981).

53. Nechaev, S. & Severinov, K. Bacteriophage-induced modifications of host RNA polymerase. Annu. Rev. Microbiol. 57, 301–322 (2003).

54. Minakhin, L. et al. Genome sequence and gene expression of *Bacillus anthracis* bacteriophage Fah. J. Mol. Biol. 354, 1–15 (2005).

55. Wörmer, L. et al. Microbial dormancy in the marine subsurface: Global endospore abundance and response to burial. Sci Adv 5, eaav1024 (2019).

56. Barylski, J. et al. Analysis of spounaviruses as a case study for the overdue reclassification of tailed phages. Syst. Biol. 69, 110–123 (2020).

57. Swanson, M. M. et al. Novel bacteriophages containing a genome of another bacteriophage within their genomes. PLoS One 7, e40683 (2012).

58. Yee, L. M. et al. The genome of *Bacillus subtilis* phage SP10: a comparative analysis with phage SPO1. Biosci., Biotechnol., Biochem. 75, 944–952 (2011).

59. Lonetto, M., Gribskov, M. & Gross, C. A. The sigma 70 family: sequence conservation and evolutionary relationships. J. Bacteriol. 174, 3843–3849 (1992).

60. Twist, K.-A. F. et al. Crystal structure of the bacteriophage T4 late-transcription coactivator gp33 with the β-subunit flap domain of *Escherichia coli* RNA polymerase. Proc. Natl. Acad. Sci. USA 108, 19961–19966 (2011).

61. Sokolova, M. et al. A non-canonical multisubunit RNA polymerase encoded by the AR9 phage recognizes the template strand of its uracil-containing promoters. Nucleic Acids Res. 45, 5958–5967 (2017).

62. Jaishankar, J. & Srivastava, P. Molecular basis of stationary phase survival and applications. Front Microbiol 8, 2000 (2017).

63. Rodriguez-Gonzalez, R. A., Leung, C. Y., Chan, B. K., Turner, P. E. & Weitz, J. S. Quantitative models of phage-antibiotic combination therapy. mSystems 5, e0075619 (2020).

64. Breitbart, M., Bonnain, C., Malki, K. & Sawaya, N. A. Phage puppet masters of the marine microbial realm. Nat Microbiol 3, 754–766 (2018).

65. Mihara, T. et al. Linking virus genomes with host taxonomy. Viruses 8, 66 (2016).

66. Eddy, S. R. Accelerated profile HMM searches. *PLoS Comp*. Biol. 7, e1002195 (2011).

67. Burton, A. T., DeLoughery, A., Li, G.-W. & Kearns, D. B. Transcriptional regulation and mechanism of SigN (ZpdN), a pBS32-encoded sigma factor in *Bacillus subtilis*. mBio 10, e0189919 (2019).

68. Katoh, K. & Standley, D. M. MAFFT multiple sequence alignment software version 7: improvements in performance and usability. Mol. Biol. Evol. 30, 772–780 (2013).

69. Capella-Gutiérrez, S., Silla-Martínez, J. M. & Gabaldón, T. trimAl: a tool for automated alignment trimming in large-scale phylogenetic analyses. Bioinformatics 25, 1972–1973 (2009).

70. Kozlov, A. M., Darriba, D., Flouri, T., Morel, B. & Stamatakis, A. RAxML-NG: a fast, scalable and user-friendly tool for maximum likelihood phylogenetic inference. Bioinformatics 35, 4453–4455 (2019).

71. Darriba, D. et al. ModelTest-NG: a new and scalable tool for the selection of DNA and protein evolutionary models. Mol. Biol. Evol. 37, 291–294 (2020).

72. Lemoine, F. et al. Renewing Felsenstein’s phylogenetic bootstrap in the era of big data. Nature 556, 452–456 (2018).

73. Xu, S. et al. ggtreeExtra: Compact visualization of richly annotated phylogenetic data. Mol. Biol. Evol. 38, 4039–4042 (2021).

74. Harwood, C. R. & Cutting, S. M. Molecular biological methods for Bacillus. (Wiley, 1990).

75. Kauffman, K. M. & Polz, M. F. Streamlining standard bacteriophage methods for higher throughput. MethodsX 5, 159–172 (2018).

76. Schilling, T., Dietrich, S., Hoppert, M. & Hertel, R. A CRISPR-Cas9-based toolkit for fast and precise in vivo genetic engineering of *Bacillus subtilis* phages. Viruses 10, 241 (2018).

77. Altenbuchner, J. Editing of the *Bacillus subtilis* genome by the CRISPR-Cas9 system. Appl. Environ. Microbiol. 82, 5421–5427 (2016).

78. Westers, H. et al. Genome engineering reveals large dispensable regions in *Bacillus subtilis*. Mol. Biol. Evol. 20, 2076–2090 (2003).

79. Storms, Z. J., Teel, M. R., Mercurio, K. & Sauvageau, D. The virulence index: a metric for quantitative analysis of phage virulence. Phage (New Rochelle) 1, 27–36 (2020).

80. Bolger, A. M., Lohse, M. & Usadel, B. Trimmomatic: a flexible trimmer for Illumina sequence data. Bioinformatics 30, 2114–2120 (2014).

81. Langmead, B. & Salzberg, S. L. Fast gapped-read alignment with Bowtie 2. Nat. Methods 9, 357–359 (2012).

82. Liao, Y., Smyth, G. K. & Shi, W. featureCounts: an efficient general purpose program for assigning sequence reads to genomic features. Bioinformatics 30, 923–930 (2014).

83. Love, M. I., Huber, W. & Anders, S. Moderated estimation of fold change and dispersion for RNA-seq data with DESeq2. Genome Biol 15, 1–21 (2014).

84. R: A Language and Environment for Statistical Computing (R Foundation for Statistical Computing, Vienna, Austria, 2021).

85. Karava, M., Bracharz, F. & Kabisch, J. Quantification and isolation of *Bacillus subtilis* spores using cell sorting and automated gating. PLoS One 14, e0219892 (2019).

86. Zhu, B. & Stülke, J. Subti Wiki in 2018: from genes and proteins to functional network annotation of the model organism *Bacillus subtilis*. Nucleic Acids Res. 46, D743–D748 (2018).

