## Supplement for "Phage-encoded sigma factors alter bacterial dormancy"

**Table S1.** Bacteria and phage strains used in this study

| | Strain | Source |
|---|---|---|
| Bacteria | *Bacillus subtilis* Δ6 | Bacillus Genetic Stock Center. BGSCID 1A1299 [1] |
| | *Bacillus subtilis* TS01 | Robert Hertel [2] |
| | *Bacillus megaterium* KM (Eldridge host) | Center for phage technology, Texas A&M University, ATCC #13632 |
| Phage | SP10 | Félix d'Hérelle Reference Center for bacterial viruses of the Université Laval [3] |
| | Goe3 | Robert Hertel [4] |
| | Eldridge | Louise Temple [5] |


1. Westers, H. *et al.* Genome engineering reveals large dispensable regions in *Bacillus subtilis*. *Mol. Biol. Evol.* **20**, 2076-2090 (2003).
2. Schilling, T., Dietrich, S., Hoppert, M. & Hertel, R. A CRISPR-Cas9-based toolkit for fast and precise in vivo genetic engineering of *Bacillus subtilis* phages. *Viruses* **10**, 241 (2018).
3. Yee, L. M. *et al.* The genome of *Bacillus subtilis* phage SP10: a comparative analysis with phage SPO1. *Biosci., Biotechnol., Biochem.* **75**, 944-952 (2011).
4. Willms, I. M., Hoppert, M. & Hertel, R. Characterization of *Bacillus subtilis* viruses vB_BsuM-Goe2 and vB_BsuM-Goe3. *Viruses* **9**, 146 (2017).
5. Reveille, A. M., Eldridge, K. A. & Temple, L. M. Complete genome sequence of *Bacillus megaterium* bacteriophage Eldridge. *Genome Announc.* **4**, e0172815 (2016).

**Table S2**. Spore yield statistics. We used flow cytometry to quantify spores and vegetative cells and calculate the fraction of spores in cultures of *Bacillus subtilis* strains with sigma factors cloned under an IPTG-inducible promoter (indicated in "group 1" column), or with empty vector controls ("group 2"). The mean value of each group is the ratio of spore fraction in induced vs. non-induced cultures (n>8). We used a two-tailed Welch two Sample *t*-test, (difference in means ≠ 0) to compare cloned gene effect on sporulation between each of the cloned genes and the empty vector controls.

| Group 1 | Group 2 | *t*-value | df | *P*-value | adjusted *P* [1] | | Mean group 1 | Mean group 2 | difference in means ± CI [2] |
|---|---|---|---|---|---|---|---|---|---|
| sigF | empty vector | 6.432 | 8.199 | 0.000182 | 0.000545 | *** | 0.1585 | 1.024 | 0.865 ± 0.309 |
| sigG | empty vector | 2.769 | 13.86 | 0.01519 | 0.02279 | * | 0.4906 | 1.024 | 0.533 ± 0.413 |
| ELDg168 | empty vector | 2.097 | 15.12 | 0.05324 | 0.06389 | | 0.6611 | 1.024 | 0.363 ± 0.368 |
| ELDg169 | empty vector | 7.802 | 7.003 | 0.000107 | 0.000545 | *** | 0.01658 | 1.024 | 1.01 ± 0.305 |
| Goe3 | empty vector | 4.387 | 8.864 | 0.001818 | 0.003635 | ** | 0.4203 | 1.024 | 0.603 ± 0.312 |
| SP10 | empty vector | 0.4636 | 10.36 | 0.6525 | 0.6525 | | 0.9567 | 1.024 | 0.067 ± 0.321 |

1. Multiple testing correction by the Benjamini, Hochberg, and Yekutieli method (* = $P < 0.05$, ** = $P < 0.01$, *** = $P < 0.001$)
2. CI = 95 % confidence interval

**Table S3.** Statistics associated with total cell densities following sigma factors induction. We used flow cytometry to quantify spores and vegetative cells in *Bacillus subtilis* strains with cloned sigma factor genes. We used a two-tailed One Sample *t*-test (mean ≠ 0) to compare the difference in cell densities between induced and non-induced paired samples.

| cloned gene | cell type | *t*-value | df | *P*-value | adjusted *P* [1] | | mean ± CI [2] |
|---|---|---|---|---|---|---|---|
| empty vector | spores | 0.3994 | 7 | 0.7015 | 0.7015 | | 3.37e+06 ± 1.99e+07 |
| sigF | spores | -4.05 | 7 | 0.00487 | 0.01136 | * | -6.62e+07 ± 3.86e+07 |
| sigG | spores | -5.491 | 7 | 0.0009151 | 0.005003 | ** | -6.28e+07 ± 2.70e+07 |
| SP10 | spores | -4.402 | 7 | 0.003148 | 0.009444 | ** | -3.18e+07 ± 1.71e+07 |
| Goe3 | spores | -5.453 | 7 | 0.000953 | 0.005003 | ** | -5.14e+07 ± 2.23e+07 |
| ELDg168 | spores | -4.114 | 9 | 0.002622 | 0.009444 | ** | -3.58e+07 ± 1.97e+07 |
| ELDg169 | spores | -13.81 | 9 | 2.30E-07 | 4.83E-06 | *** | -8.80e+07 ± 1.44e+07 |
| empty vector | vegetative | 1.115 | 7 | 0.3015 | 0.3333 | | 1.10e+07 ± 2.33e+07 |
| sigF | vegetative | 0.7883 | 7 | 0.4564 | 0.4792 | | 2.09e+07 ± 6.28e+07 |
| sigG | vegetative | -2.341 | 7 | 0.05178 | 0.09062 | | -2.51e+07 ± 2.53e+07 |
| SP10 | vegetative | -2.981 | 7 | 0.0205 | 0.03913 | * | -1.47e+07 ± 1.17e+07 |
| Goe3 | vegetative | 1.569 | 7 | 0.1607 | 0.2249 | | 2.35e+07 ± 3.54e+07 |
| ELDg168 | vegetative | 1.768 | 9 | 0.1108 | 0.179 | | 1.76e+07 ± 2.25e+07 |
| ELDg169 | vegetative | 1.368 | 9 | 0.2045 | 0.2526 | | 9.13e+06 ± 1.51e+07 |
| empty vector | total cells | 1.49 | 7 | 0.1797 | 0.2359 | | 1.44e+07 ± 2.28e+07 |
| sigF | total cells | -1.191 | 7 | 0.2723 | 0.3177 | | -4.52e+07 ± 8.97e+07 |
| sigG | total cells | -4.109 | 7 | 0.004523 | 0.01136 | * | -8.79e+07 ± 5.06e+07 |
| SP10 | total cells | -4.522 | 7 | 0.002727 | 0.009444 | ** | -4.66e+07 ± 2.44e+07 |
| Goe3 | total cells | -1.599 | 7 | 0.1539 | 0.2249 | | -2.79e+07 ± 4.12e+07 |
| ELDg168 | total cells | -2.896 | 9 | 0.0177 | 0.03718 | * | -1.82e+07 ± 1.42e+07 |
| ELDg169 | total cells | -8.405 | 9 | 1.49E-05 | 0.0001563 | *** | -7.89e+07 ± 2.12e+07 |

1. Multiple testing correction by the Benjamini, Hochberg, and Yekutieli method (* = $P < 0.05$, ** = $P < 0.01$, *** = $P < 0.001$)
2. For each paired samples the difference in cell densities (per mL) are calculated as *induced culture – non-induced culture.* CI = 95 % confidence interval

**Table S4.** Viral orthologous groups (VOG) of sigma factors

| Group Name | Consensus Functional Description | Protein Count | Species Count |
|---|---|---|---|
| VOG00048 | sp\|P33658\|RPSG_CLOAB RNA polymerase sigma-G factor | 261 | 189 |
| VOG05673 | sp\|P04524\|RP55_BPT4 RNA polymerase sigma factor | 286 | 286 |
| VOG07471 | sp\|P03048\|RP28_BPSP1 RNA polymerase sigma GP28 factor | 6 | 6 |
| VOG12906 | REFSEQ RNA polymerase sigma factor | 4 | 4 |
| VOG16626 | REFSEQ RNA polymerase sigma factor | 7 | 7 |
| VOG18372 | REFSEQ sigma70, RNA polymerase sigma factor, positive control factor Xpf (N-terminal region) | 2 | 2 |
| VOG20983 | REFSEQ sigma-70 family RNA polymerase sigma factor | 2 | 2 |
| VOG26037 | REFSEQ RNA polymerase sigma factor SigF | 2 | 2 |

**Table S5.** TIGRFAM* protein families of bacterial sigma factors

| ID | Accession | Description | Label in main Fig. 2 |
|---|---|---|---|
| rpoH_proteo | TIGR02392 | alternative sigma factor RpoH | other |
| RpoD_Cterm | TIGR02393 | RNA polymerase sigma factor RpoD | other |
| rpoS_proteo | TIGR02394 | RNA polymerase sigma factor RpoS | other |
| rpoN_sigma | TIGR02395 | RNA polymerase sigma-54 factor | other |
| FliA_WhiG | TIGR02479 | RNA polymerase sigma factor, FliA/WhiG family | other |
| spore_sigmaE | TIGR02835 | RNA polymerase sigma-E factor | sigE |
| spore_sigmaK | TIGR02846 | RNA polymerase sigma-K factor | sigK |
| spore_sigG | TIGR02850 | RNA polymerase sigma-G factor | sigG |
| spore_sigH | TIGR02859 | RNA polymerase sigma-H factor | other |
| spore_sigF | TIGR02885 | RNA polymerase sigma-F factor | sigF |
| spore_sigI | TIGR02895 | RNA polymerase sigma-I factor | other |
| sigma70-ECF | TIGR02937 | RNA polymerase sigma factor, sigma-70 family | other |
| RpoE_Sigma70 | TIGR02939 | RNA polymerase sigma factor RpoE | other |
| Sigma_B | TIGR02941 | RNA polymerase sigma-B factor | other |
| Sig70_famx1 | TIGR02943 | RNA polymerase sigma-70 factor, TIGR02943 family | other |
| SigH_actino | TIGR02947 | RNA polymerase sigma-70 factor, TIGR02947 family | other |
| SigW_bacill | TIGR02948 | RNA polymerase sigma-W factor | other |
| SigM_subfam | TIGR02950 | RNA polymerase sigma factor, SigM family | other |
| Sig70_famx2 | TIGR02952 | RNA polymerase sigma-70 factor, TIGR02952 family | other |
| Sig70_famx3 | TIGR02954 | RNA polymerase sigma-70 factor, TIGR02954 family | other |
| SigX4 | TIGR02957 | RNA polymerase sigma-70 factor, TIGR02957 family | other |
| SigZ | TIGR02959 | RNA polymerase sigma factor, SigZ family | other |
| SigX5 | TIGR02960 | RNA polymerase sigma-70 factor, TIGR02960 family | other |
| SigBFG | TIGR02980 | RNA polymerase sigma-70 factor, sigma-B/F/G subfamily | other |
| SigE-fam_strep | TIGR02983 | RNA polymerase sigma-70 factor, sigma-E family | other |
| Sig-70_plancto1 | TIGR02984 | RNA polymerase sigma-70 factor, Planctomycetaceae-specific subfamily 1 | other |
| Sig70_bacteroi1 | TIGR02985 | RNA polymerase sigma-70 factor, Bacteroides expansion family 1 | other |
| Sig-70_gvs1 | TIGR02989 | RNA polymerase sigma-70 factor, Rhodopirellula/Verrucomicrobium family | other |
| Sig70-cyanoRpoD | TIGR02997 | RNA polymerase sigma factor, cyanobacterial RpoD-like family | other |
| Sig-70_X6 | TIGR02999 | RNA polymerase sigma factor, TIGR02999 family | other |
| Sig-70_gmx1 | TIGR03001 | RNA polymerase sigma-70 factor, Myxococcales family 1 | other |

* ncbi.nlm.nih.gov/hmm/TIGRFAMs/release_15.0

**Table S6.** Plasmids used in this study

| name | Backbone / RE digest | Insert amplicon / RE digest | source |
| --- | --- | --- | --- |
| pJOE88889 | - | - | BGSC ECE358 |
| pSP10-sgRNA | pJOE88889 | SP10_KOsigF_1+2 / BsaI | This study |
| pSP10-delete120 | pSP10-sgRNA | SP10_KOsigF_3+4+5+6 / SfiI | This study |
| pDR110 | - | - | David Rudner, Xindan Wang |
| pDAS1 | pDR110/NheI + SphI | oDAS1+oDAS2 /XbaI + SphI | This study |
| pDAS2 | pDR110/NheI + SphI | oDAS3+oDAS4 /XbaI + SphI | This study |
| pDAS3 | pDR110/NheI + SphI | oDAS5+oDAS5 /XbaI + SphI | This study |
| pDAS4 | pDR110/NheI + SphI | oDAS7+oDAS8 /XbaI + SphI | This study |
| pDAS5 | pDR110/NheI + HindIII | oDAS1+oDAS2 /XbaI + HindIII | This study |
| pDAS6 | pDR110/NheI + HindIII | oDAS1+oDAS2 /XbaI + HindIII | This study |

**Table S7.** Primers used in this study

| Name | target | Sequence* |
|---|---|---|
| SP10_KOsigF_1 | SP10 (sgRNA) | tacgGATCATACCTGTTGTAAGCG |
| SP10_KOsigF_2 | SP10 (sgRNA) | aaacCGCTTACAACAGGTATGATC |
| SP10_KOsigF_3 | SP10 (g120 flank) | taggatccggccaacgaggccCAAAAAGTTGTAATCGGTATTGACTGGGG |
| SP10_KOsigF_4 | SP10 (g120 flank) | aagtagtccactaCATAACTAGACCTCCCGTTGTTTTCC |
| SP10_KOsigF_5 | SP10 (g120 flank) | aggtctagttatgTAGTGGACTACTTCAAAACCTACGAAGG |
| SP10_KOsigF_6 | SP10 (g120 flank) | taggatccggccttattggccCTCGACTTGACTGTTATCTTTAGAAACAATGC |
| SP10_valid_F | SP10 (g120 flank) | CCCTAAAACCTCCCGAGCTG |
| SP10_valid_R | SP10 (g120 flank) | GTCTTCCATGATACCGCCCC |
| oDAS1 | SP10-F | ctctctagaacataaggaggaactactATGTCTAAGAATTTTAATCCAGC |
| oDAS2 | SP10-R | ctcgcatgcCTATTGTATCACCTCTTTTTGG |
| oDAS3 | Goe3-F | ctctctagaacataaggaggaactactATGGGTAAGATTAAAACC |
| oDAS4 | Goe3-R | ctcgcatgcTTAGACATTGGTTTTACCCTCC |
| oDAS5 | 168SigF-F | ctctctagaacataaggaggaactactATGGATGTGGAGGTTAAGAAAAACG |
| oDAS6 | 168SigF-R | ctcgcatgcCTAGCCATCCGTATGATCCATTTG |
| oDAS7 | 168SigG-F | ctctctagaacataaggaggaactactGTGTCGAGAAATAAAGTCGAAATC |
| oDAS8 | 168SigG-R | ctcgcatgcTTATTGATGAATATTTTTATTCATTTGTTTGATAG |
| oDAS9 | pDR110 (screen) | CGTACGATCTTTCAGCCG |
| oDAS10 | pDR110 (screen) | AGAACGTTGCTCGAGGG |
| oDAS15 | Eldridge169-F | ctcaagcttacataaggaggaactactGTGGTGGGTAAGAGCAAATATC |
| oDAS16 | Eldridge169-R | ctctctagaTTAGCTTCCCACGTTAATACGG |
| oDAS17 | Eldridge168-F | ctcaagcttacataaggaggaactactATGAGTAAAAAAGAATTCT |
| oDAS18 | Eldridge168-R | ctctctagaTTACTGAGCAGCTACTAG |

* Target binding nucleotides are in uppercase letters and 5’ overhangs are in lower case letters. Added restriction sites are underlined.

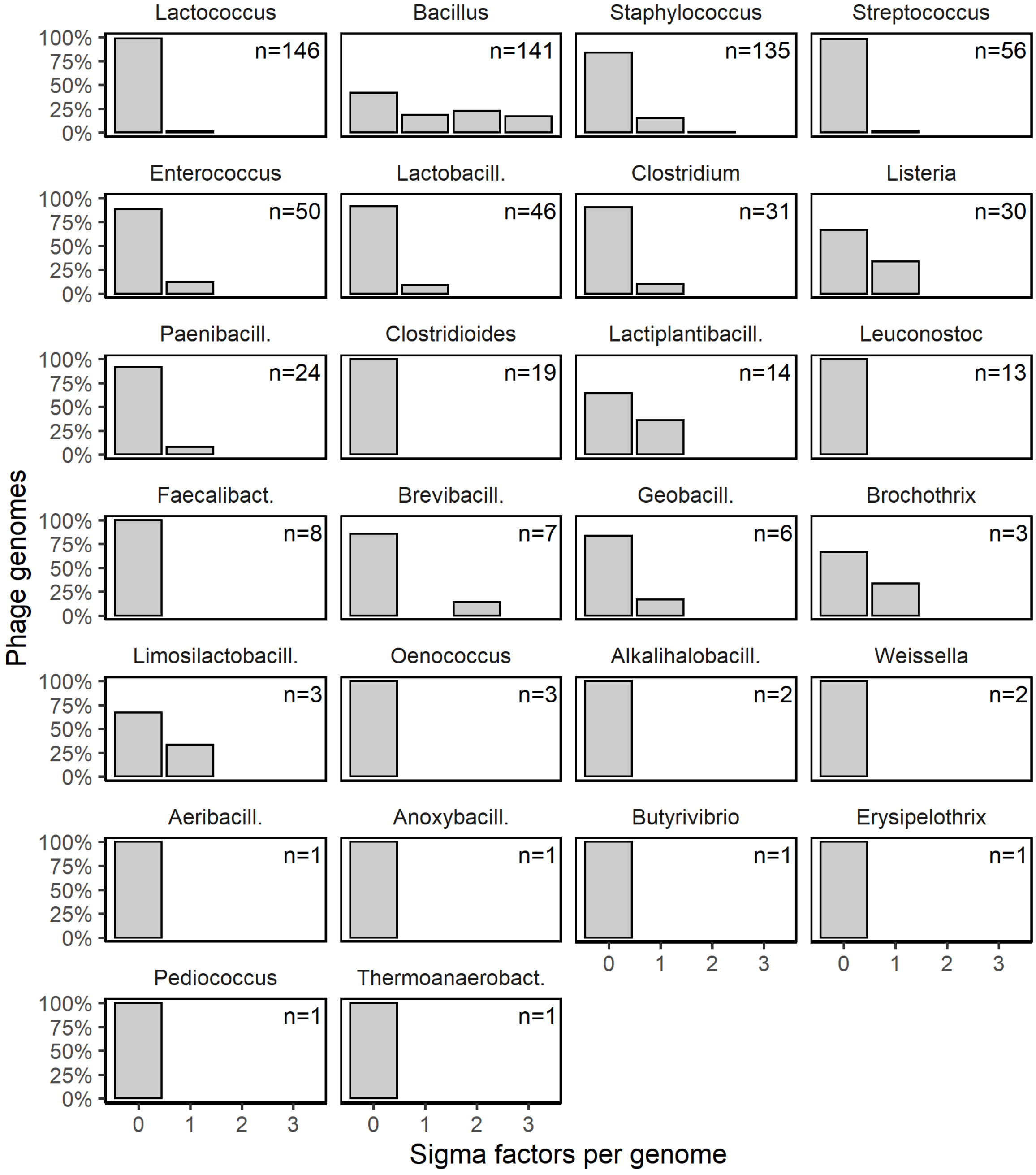


**Fig. S1.** Multiplicity of sigma factors in the genomes of phages that infect different *Firmicutes* genera. Each facet shows the distribution of sigma factor gene counts in genomes of phages infecting a single genus. Sigma factor genes were obtained from the Virus Orthologous Database and mapped to hosts using the Virus-Host Database.

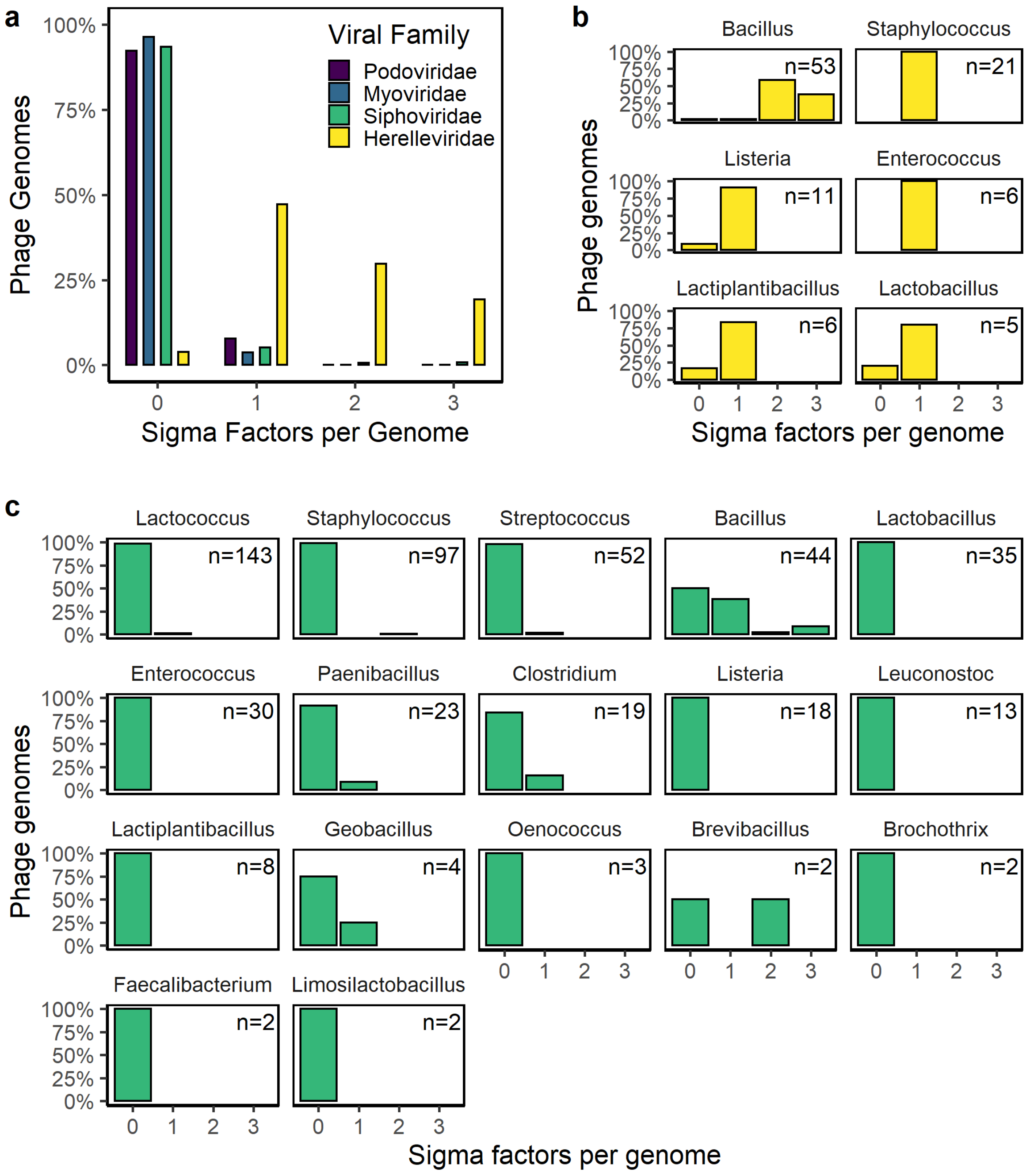


**Fig. S2.** Multiplicity of sigma factors in the genomes of phages that in infect *Firmicutes* by viral family. **a,** Percent of phage genomes containing different numbers of sigma-factor genes is shown across phage families that infect *Firmicutes* (*Podoviridae*, n = 77; *Myoviridae*, n = 55; *Siphoviridae*, n = 503; *Herelleviridae*, n = 104). **b-c**, The data from **a** is separated out into panels by host genera for two viral families having members encoding multiple sigma factors: *Herelleviridae* (**b**) and *Siphoviridae* (**c**). Sigma-factor genes were obtained from the Virus Orthologous Database and mapped to hosts using the Virus-Host Database.

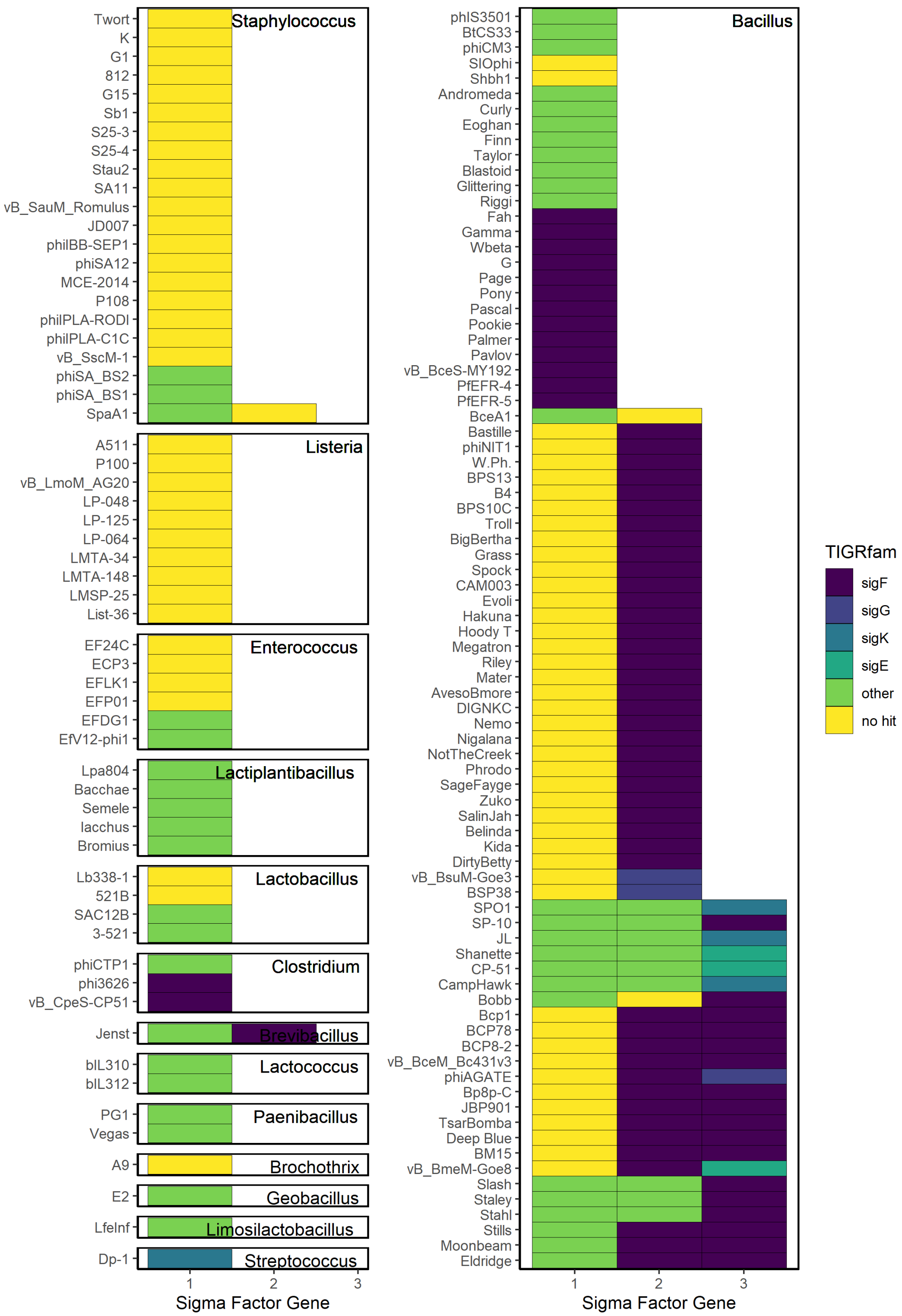


**Fig. S3.** Sigma factor composition in *Firmicutes* phages. Host genera are depicted with black text labels in the upper right of each panel. Each row shows the type of sigma factors in a single phage genome. Each phage-encoded sigma factor gene was assigned to a bacterial sigma-factor family based on best hit (by sequence E-value) when queried against a collection of bacterial-encoded sigma-factor protein profiles from TIGRFAM using hmmscan.

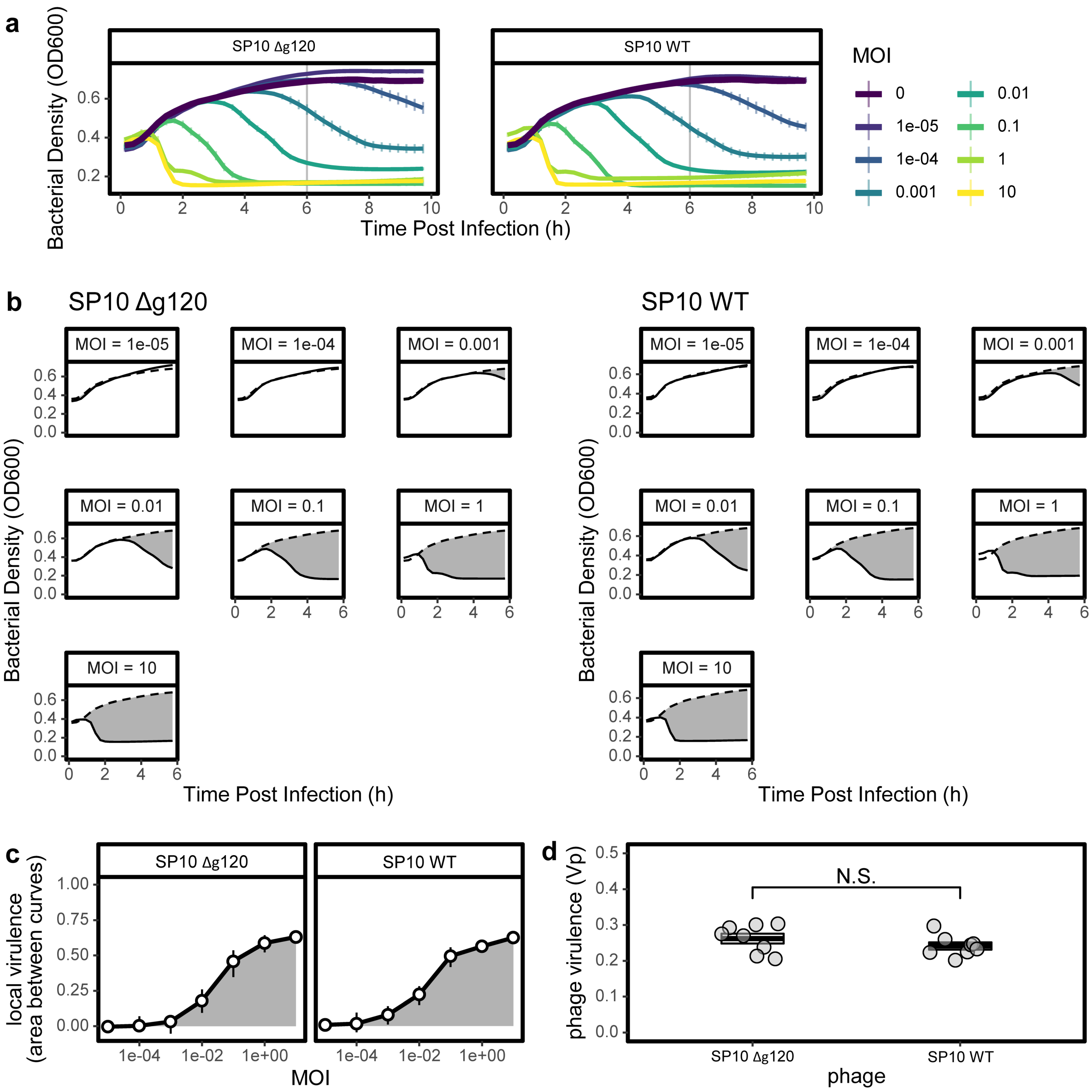


**Fig. S4. A sporulation-specific sigma factor is non-essential for phage virulence.** Phage SP10 from which we deleted the sporulation sigma factor homolog g120 (Δg120) is as virulent as the wild-type (WT) phage when infecting *Bacillus subtilis* Δ6. **a-d**, For measuring phage virulence (Vp) (**a**) we measured optical density (OD600) to follow bacterial growth and lysis by phage over a range of multiplicities of infection (MOI = phage/bacteria titers). We used 6 h after infection (vertical grey line) as the integration limit in the next steps based on the time of transition from exponential growth to stationary phase in non-infected cultures. (**b**) from data in **a**, the MOI-specific virulence was calculated from the phage-induced reduction in bacterial growth, which is the grey area between the curves of infected (solid line) and non-infected (dashed line) cultures, as a fraction of the total area under the non-infected curve. (**c**) phage virulence was calculated from the area under the curve of local virulence vs. MOI. (**d**) The difference in virulence between phages is non-significant (N.S.) when compared using a Welch Two Sample *t*-test ($t_{12.9} = 1.22$, $P = 0.25$; n = 8).

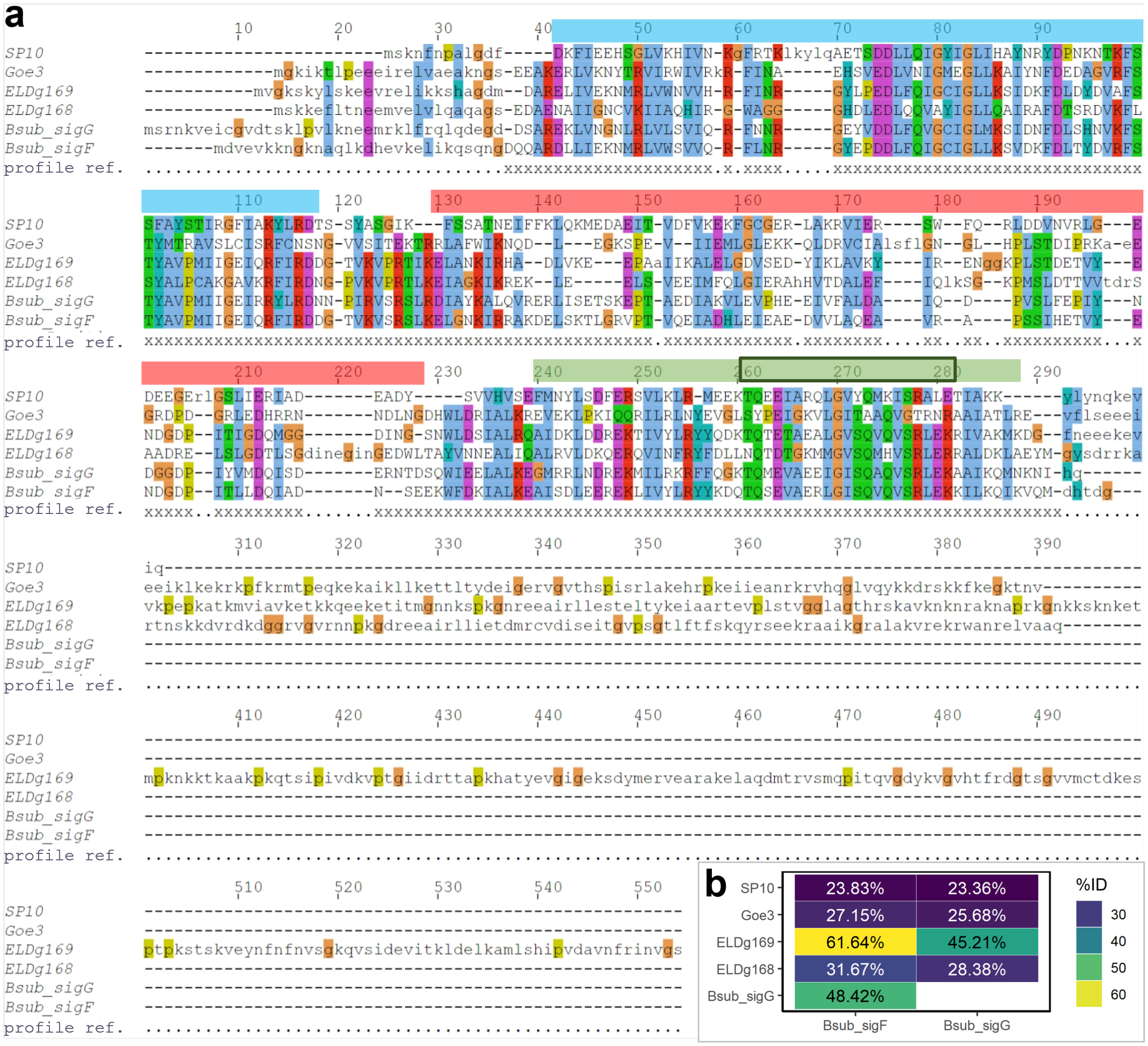


**Fig S5**. **a**, Multiple sequence alignment of sigma factors cloned in this study. Protein sequences from phages and from *Bacillus subtilis* (Bsub) were aligned to the protein profile of the RNA polymerase sigma-70 factor, sigma-B/F/G subfamily (TIGR02980) using hmmalign (HMMER v3.3). The profile reference positions are marked with ‘x’ at the bottom of the alignment. Functional sequence regions of the host sigma factors are depicted above the alignment by color bars: region 2 in blue, region 3 in red and region 4 in green, with the helix-turn-helix motif marked by a black box. Functional region annotation from SWISS-MODEL data for *B. subtilis* genes *sigF* (P07860) and *sigG* (P19940). **b**, Protein identities from the alignment. The identities were calculated using esl-alipid (part of the Easel library of HMMER v3.3) after trimming the N and C termini beyond the profile reference positions (using hmmalign –trim).

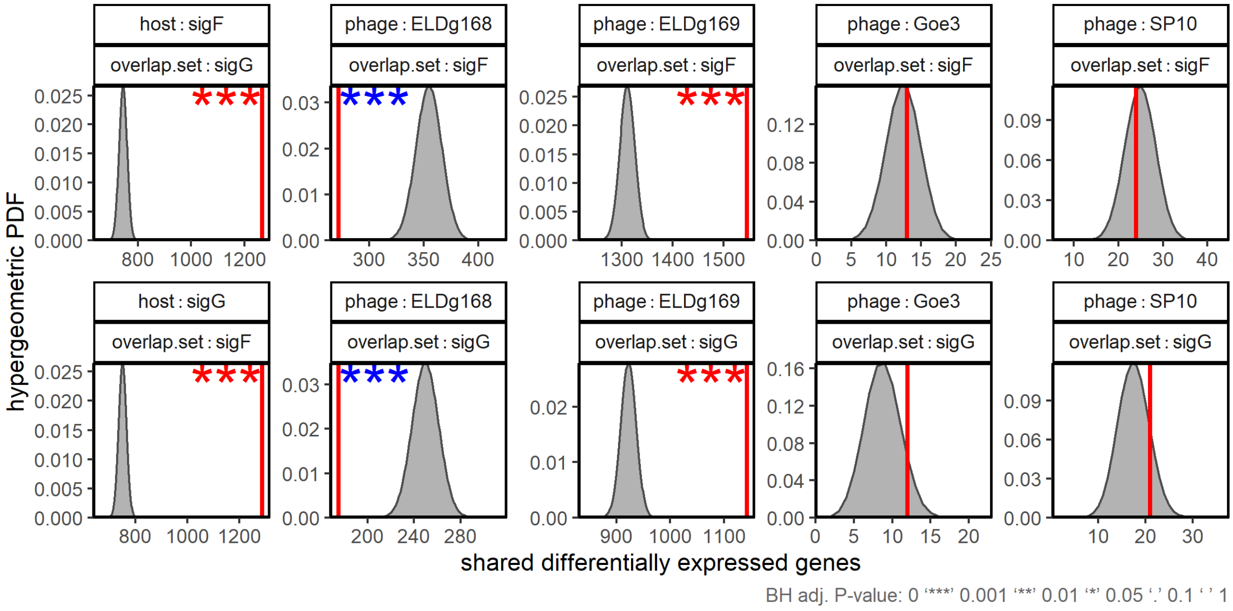


**Fig S6.** Overlap between differentially expressed genes (DEGs) following induction of phage- and bacterial-derived sigma factors. Observed (red line) versus expected (gray curve) number of shared DEGs in *Bacillus subtilis* between cultures induced to express sigma factors derived from phage (ELDg168, ELDg169, Goe3, SP10) and bacteria (*sigF*, *sigG*). Differential expression was calculated by comparing RNA-seq data from split cultures (n = 3) of induced and non-induced cells after cloning sigma factors under an IPTG inducible promoter. DEGs were considered to overlap if differential expression was significant (adjusted *P*-value < 0.05) and in the same direction (up- or down- regulated) in both sets. Asterisks indicate hypergeometric test *P*-values for the DEG overlap between the two sets being greater (red) or smaller (blue) than the random expectation. *P*-values were adjusted for multiple comparisons using Benjamini-Hochberg procedures (*** = $P < 0.001$). The random expectation is described by the probability density function (PDF) of the hypergeometric distribution given the number of DEGs in the test.

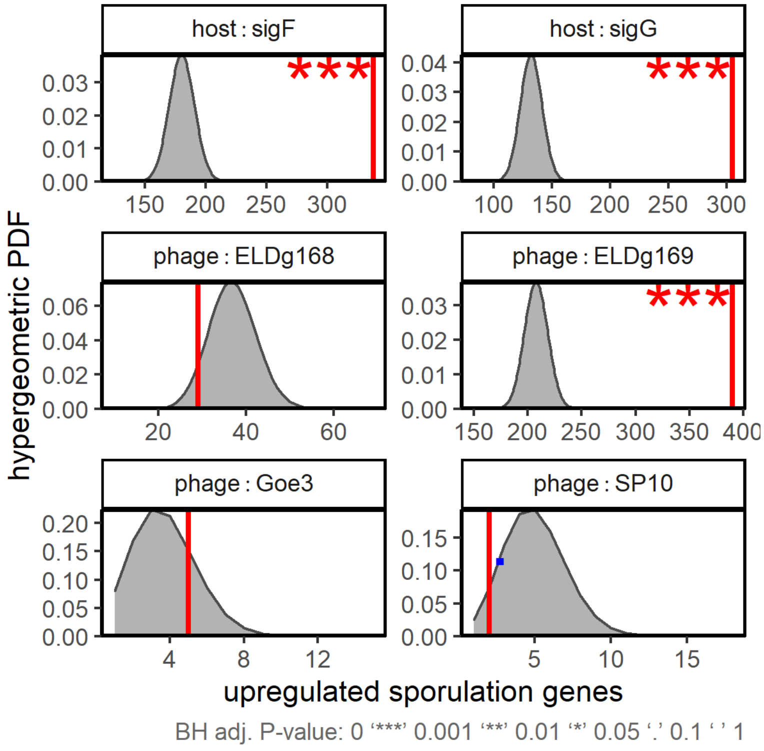


**Fig. S7.** Enrichment of upregulated sporulation genes following sigma factor induction. Observed (red line) versus expected (gray curve) number of upregulated sporulation genes in *Bacillus subtilis* cultures induced to express a sigma factor cloned under an IPTG-inducible promoter versus non-induced control cultures (n = 3). Asterisks indicate hypergeometric test *P*-values for observing more sporulation genes among upregulated genes than expected by random. *P*-values were adjusted for multiple comparisons using Benjamini-Hochberg procedures (*** = $P < 0.001$). The random expectation is described by the probability density function (PDF) of the hypergeometric distribution for 645 sporulation genes of 3,885 genes in total, and accounting for the number of observed sporulation genes among all upregulated genes for each sigma factor. Genes were considered to be upregulated when adjusted *P*-value < 0.05 and the fold change > 2. Sporulation gene categorization came from SubtiWiki.

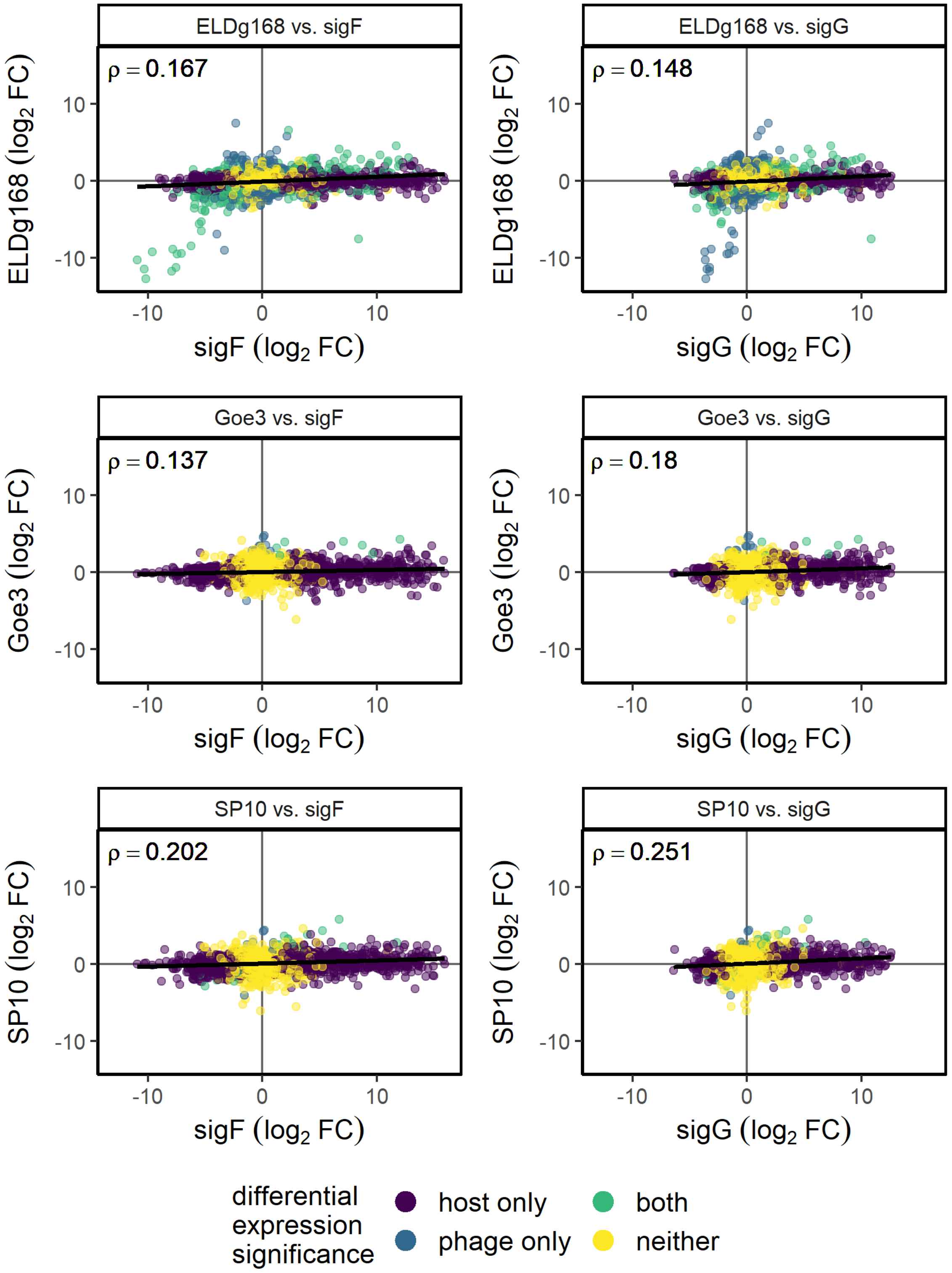


**Fig. S8.** Lack of correlation between differential gene expression when inducing the expression of phage-derived sigma factors (y-axis) and host-derived sigma factors (x-axis). See main figure 3a for details. FC = Fold Change.

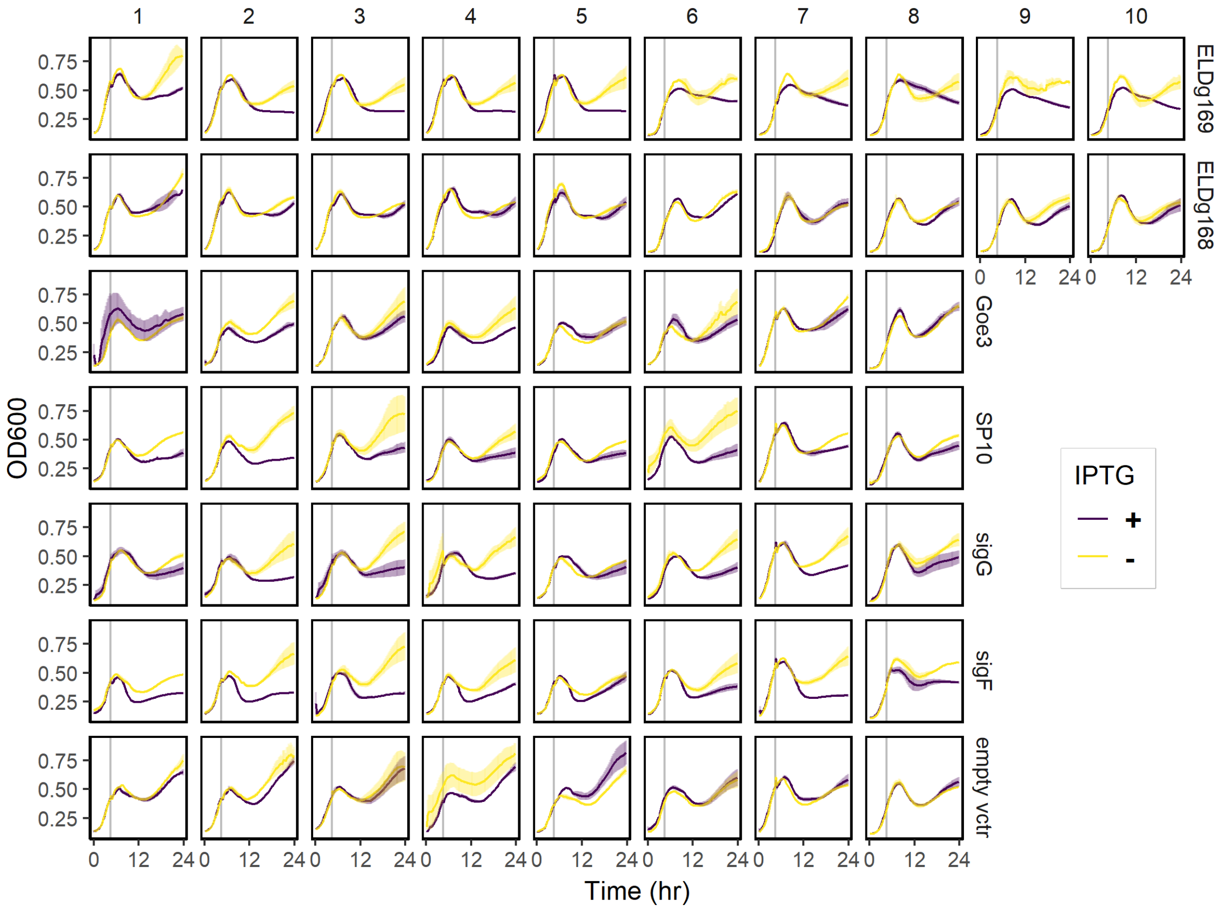


**Fig. S9.** Growth of *Bacillus subtilis* during sporulation assays. Each panel shows the growth (mean ± SE, n = 3) measured as optical density (OD600) over time of cultures in which a sigma factor (row label on the right) was cloned under an IPTG-inducible promoter. The empty-vector negative control strain had the IPTG-inducible promoter without any sigma factor. Column numbers represent independent colonies from which experimental cultures were inoculated. Cultures were grown for 4.5 h (grey vertical line) in sporulation media before IPTG was added to half of them. Samples for quantifying spores and vegetative cells by flow cytometry were taken at 24 h.

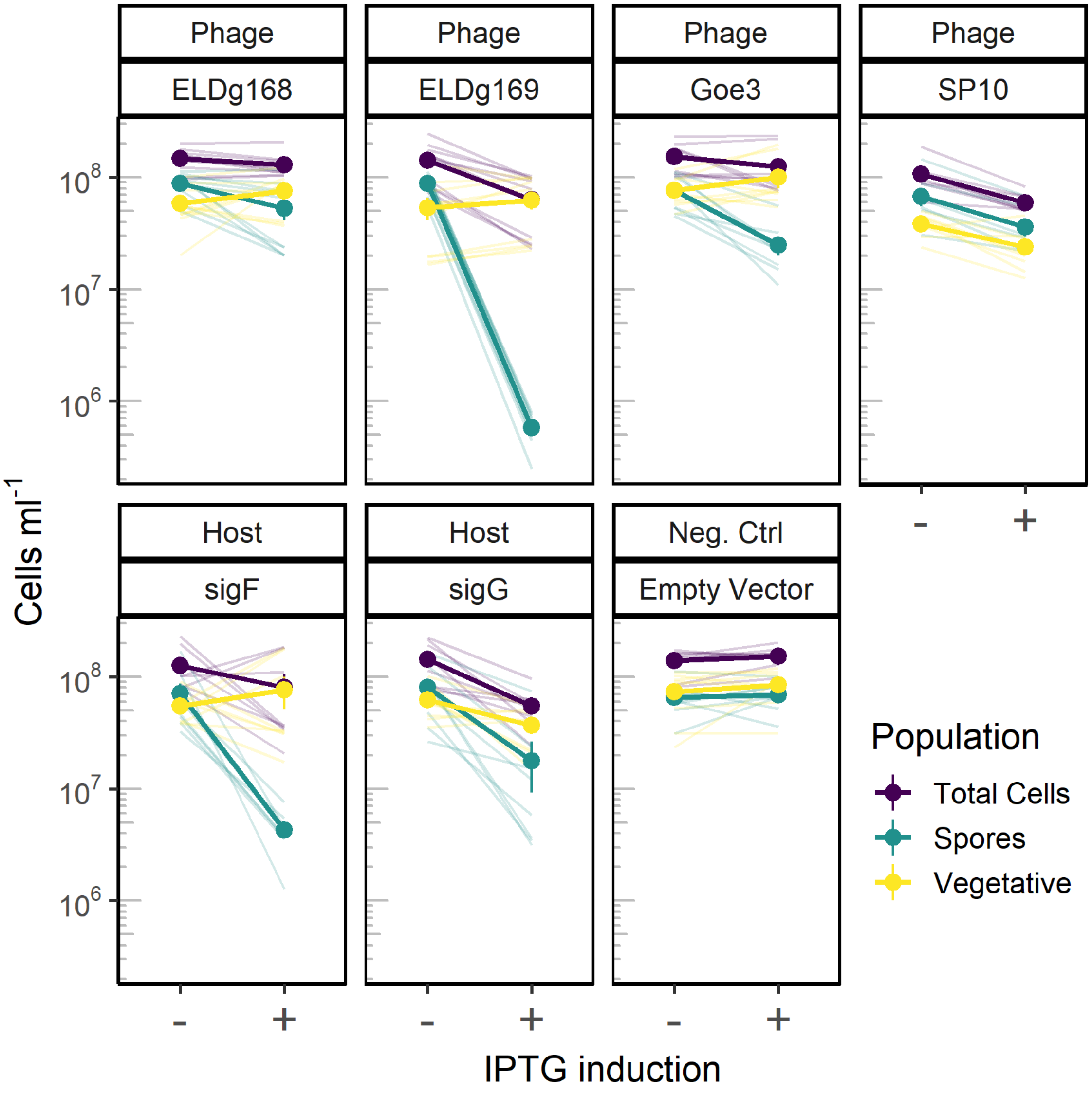


**Fig. S10.** Effect of sigma factor induction on cell density. Sporulation was measured in strains with phage-derived or bacteria-derived (host) sigma factors under an IPTG- inducible promoter. The empty-vector negative control had the IPTG-inducible promoter without any sigma factor. For each data point replicate cultures (n = 6) were grown in sporulation media for 4.5 h, before expression of the sigma factor (panel header label) was induced by adding IPTG to half the cultures. Cells were quantified and classified by flow cytometry after 24 h (spore + veg = total cells). Points connected by thick lines represent mean ± SEM of independent clones (n ≥ 8). Thin lines connect values of each induced clone and its non-induced paired control.

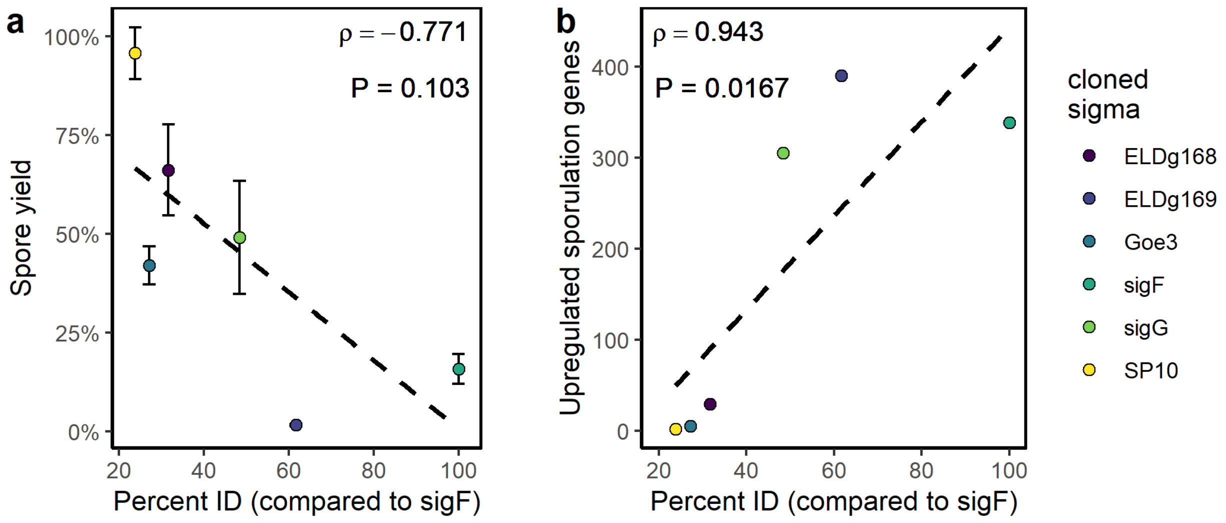


**Fig S11**. Sporulation response corresponds with the identity of sigma factors induced in *Bacillus. subtilis*. **a-b**, The amino acid identity between cloned sigma factors and *B. subtilis sigF* is negatively correlated with the mean spore yield (**a**) (see Fig. 4) when induced during sporulation and positively corelated with the number of upregulated sporulation genes (**b**) when induced in exponential phase (see Fig. 3). Spearman's rho (ρ) and the associated P-value (*P*), and linear model (dashed line) were calculated in R. Error bars in **a** represent SEM.
